# SPIDR: a highly multiplexed method for mapping RNA-protein interactions uncovers a potential mechanism for selective translational suppression upon cellular stress

**DOI:** 10.1101/2023.06.05.543769

**Authors:** Erica Wolin, Jimmy K. Guo, Mario R. Blanco, Andrew A. Perez, Isabel N. Goronzy, Ahmed A. Abdou, Darvesh Gorhe, Mitchell Guttman, Marko Jovanovic

**Author notes:** These authors contributed equally (Erica Wolin, Jimmy K. Guo). To whom correspondence should be addressed &.

## Abstract

RNA binding proteins (RBPs) play crucial roles in regulating every stage of the mRNA life cycle and mediating non-coding RNA functions. Despite their importance, the specific roles of most RBPs remain unexplored because we do not know what specific RNAs most RBPs bind. Current methods, such as crosslinking and immunoprecipitation followed by sequencing (CLIP-seq), have expanded our knowledge of RBP-RNA interactions but are generally limited by their ability to map only one RBP at a time. To address this limitation, we developed SPIDR (Split and Pool Identification of RBP targets), a massively multiplexed method to simultaneously profile global RNA binding sites of dozens to hundreds of RBPs in a single experiment. SPIDR employs split-pool barcoding coupled with antibody-bead barcoding to increase the throughput of current CLIP methods by two orders of magnitude. SPIDR reliably identifies precise, single-nucleotide RNA binding sites for diverse classes of RBPs simultaneously. Using SPIDR, we explored changes in RBP binding upon mTOR inhibition and identified that 4EBP1 acts as a dynamic RBP that selectively binds to 5’-untranslated regions of specific translationally repressed mRNAs only upon mTOR inhibition. This observation provides a potential mechanism to explain the specificity of translational regulation controlled by mTOR signaling. SPIDR has the potential to revolutionize our understanding of RNA biology and both transcriptional and post-transcriptional gene regulation by enabling rapid, *de novo* discovery of RNA-protein interactions at an unprecedented scale.

## INTRODUCTION

RNA binding proteins (RBPs) play key roles in controlling all stages of the mRNA life cycle, including transcription, processing, nuclear export, translation, and degradation^1–5^. Recent estimates suggest that up to 30% of all human proteins (several thousand in total) bind to RNA^6–10^, indicative of their broad activity and central importance in cell biology. Moreover, mutations in RBPs have been causally linked to various human diseases, including immunoregulatory and neurological disorders as well as cancer^2–4,11^. Yet, we still do not know what specific roles most of these RBPs play because the RNAs they bind remain mostly unknown.

In addition, there are many thousands of regulatory non-coding RNAs (ncRNAs) whose functional roles remain largely unknown^12, 13^; understanding how they work requires defining the proteins to which they bind^13–15^. For example, uncovering the mechanism by which the Xist long noncoding RNA (lncRNA) silences the inactive X chromosome required identification of the SPEN/SHARP RBP that binds to Xist^16–20^ – a process that took >25 years after the lncRNA was discovered^14^. Given the large discrepancy between the number of ncRNAs and putative RBPs identified, and the number of RNA-protein interactions demonstrated to be functionally relevant, there is an urgent need to generate high-resolution binding maps to enable functional characterization^14^.

Currently, the most rigorous and widely utilized method to characterize RBP-RNA interactions is crosslinking and immunoprecipitation followed by next generation sequencing (CLIP-seq)^21–26^. Briefly, CLIP works by utilizing UV light to covalently crosslink RNA and directly interacting proteins, followed by cell lysis, immunoprecipitation under stringent conditions (e.g., 1M salt) to purify a protein of interest followed by gel electrophoresis, transfer to a nitrocellulose membrane, and excision of the protein-RNA complex prior to sequencing and identification of the bound RNAs. CLIP and its related variants have greatly expanded our knowledge of RNA-RBP interactions and our understanding of gene expression from mRNA splicing to microRNA targeting^21–26^.

Yet, CLIP and all of its variants (with one recent exception^27^ which we discuss in more detail below; see **Note 1**) are limited to mapping a single RBP at a time. As such, efforts to generate reference maps for hundreds of RBPs in even a limited number of cell types have required major financial investment and the work of large teams working in international consortiums (e.g., ENCODE)^23, 28, 29^. Despite these herculean efforts and the important advances they have enabled, there are critical limitations: (i) Only a small fraction of the total number of predicted RBPs have been successfully mapped using genome-wide methods (ENCODE has so far characterized the binding patterns of < 10% of known RBPs); (ii) Of these, most have been mapped in only a small number of cell lines (mainly K562 and HepG2); (iii) Because each protein map is generated from an individual experiment, a large number of cells is required to map dozens, let alone hundreds, of RBPs – this is particularly challenging for studying primary cells, disease models, or other populations of rare cells. Further, because these datasets are highly cell type-specific, the generated maps are not likely to be directly useful for studying these RBPs within other cell-types or model systems (e.g., patient samples, animal models, or perturbations). Thus, it is critically important to enable the generation of comprehensive RBP binding for any cell type of interest in a manner that is accessible to any individual lab.

To overcome these challenges, we developed SPIDR (Split and Pool Identification of RBP targets), a massively multiplexed method to simultaneously profile the global RNA binding sites of dozens to hundreds of RBPs in a single experiment. SPIDR is based on our split-pool barcoding strategy that maps multiway nucleic acid interactions using high throughput sequencing^30–32^; the vastly simplified version of split-pool barcoding we present here, when combined with antibody-bead barcoding, increases the throughput of current CLIP methods by two orders of magnitude. Using this approach, we can reliably identify the precise, single nucleotide RNA binding sites of dozens of RBPs simultaneously and can detect changes in RBP binding upon perturbation. Using this approach, we uncovered a mechanism driven by dynamic RBP binding to mRNA that may explain the specificity of translational regulation controlled by mTOR signaling. Thus, SPIDR enables rapid, *de novo* discovery of RNA-protein interactions at an unprecedented scale and has the potential to transform our understanding of RNA biology and both transcriptional and post-transcriptional gene regulation.

## RESULTS

### SPIDR: A highly multiplexed method for mapping RBP-RNA interactions

We developed SPIDR to enable highly multiplexed mapping of RBPs to individual RNAs transcriptome-wide. Briefly, SPIDR involves: (i) generating highly multiplexed antibody-bead pools by tagging individual antibody-bead conjugates with a specific oligonucleotide (tagged bead pools), (ii) performing RBP purification using these tagged antibody-bead pools in UV-crosslinked cell lysates, and (iii) linking individual antibodies to their associated RNAs using split-and-pool barcoding (**Figure 1A** and **Supplemental Figure 1**).

**Figure 1:**
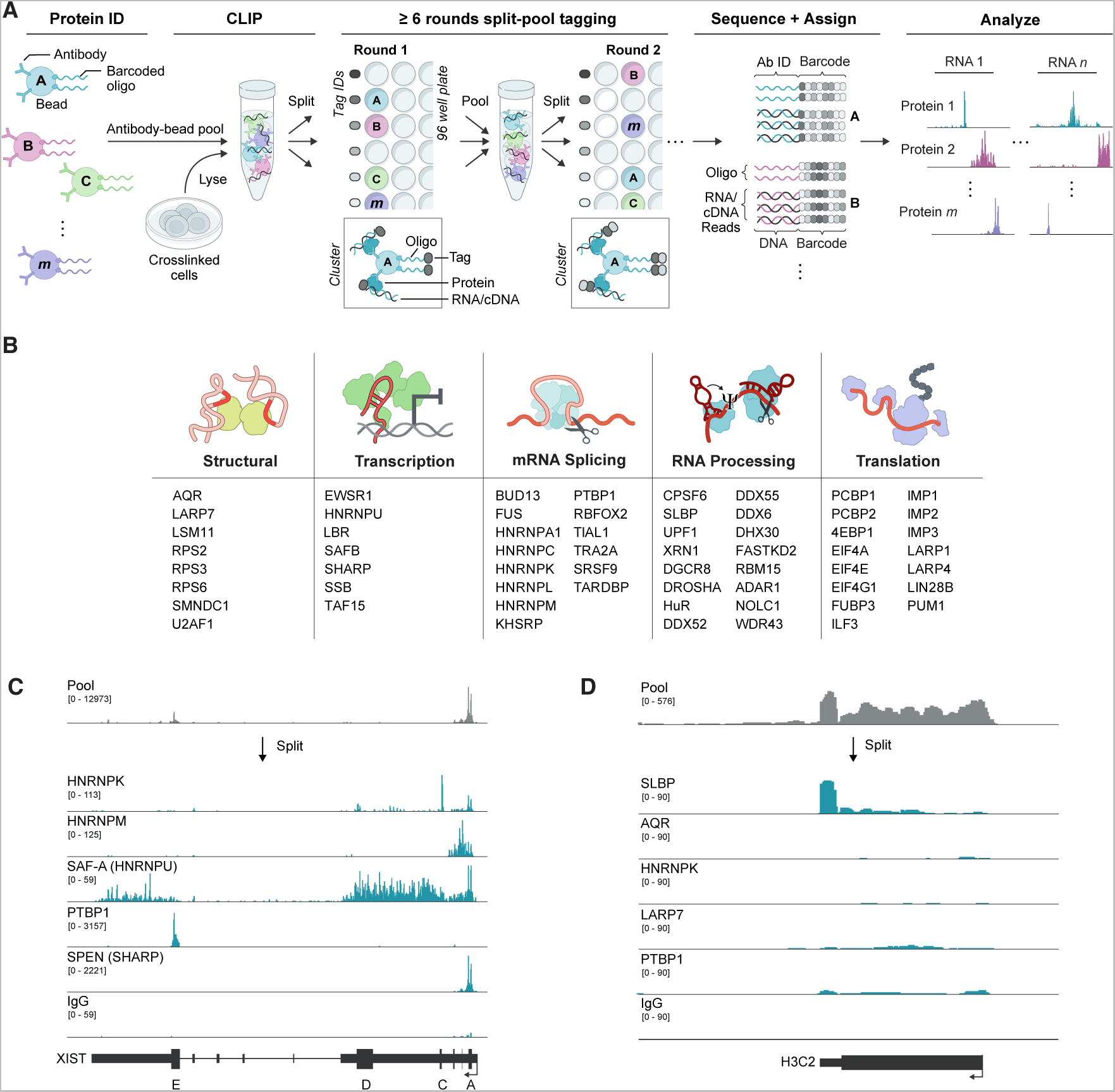
SPIDR (Split and Pool Identification of RBP targets) – a highly multiplexed method to map protein-RNA interactions. **(A)** Schematic overview of the SPIDR method. The bead pool is incubated with UV crosslinked lysate in a single tube. After immunopurification, each bead is uniquely labeled by split-and-pool barcoding. The complexity of the barcode generated depends on the number of individual tags used in each split-pool round and the number of split-pool rounds. For example, after 8 rounds of split and pool barcoding, using 12 barcodes in each round, the likelihood that two beads will end up with the same barcode is ~ 1 in 430 million (1/128). Oligos and RNA molecules and their linked barcodes are sequenced and RNAs are matched to proteins based on their shared barcodes. (The bead labeling strategy was adapted from ChIP-DIP, a protocol for multiplexed mapping of proteins to DNA, https://guttmanlab.caltech.edu/ <u>technologies/</u>). **(B)** Schematic list of the different RBPs mapped by SPIDR in K562 and/or HEK293T cells, functional assignments based on literature review. **(C)** An example of the raw alignment data for the pool (all reads before splitting by bead identities) and for specific RBPs (all reads assigned to specific RBP beads) across the *XIST* RNA. Blocks represent exons, lines introns, and thick blocks are the annotated *XIST* repeat regions (A-E). **(D)** Raw alignment data for SLBP across the *H3C2* histone mRNA. Top track is pooled alignment data; tracks below are reads assigned to SLBP or other RBPs and controls.

We first devised a highly modular scheme to generate hundreds of tagged antibody-beads such that each unique bead population is labeled with a specific oligonucleotide tag and all bead populations are combined to generate an antibody-bead pool (**Figure 1A** and **Supplemental Figure 1**). Because this approach does not require direct chemical modification of the antibody, we can utilize any antibody (in any storage buffer) and rapidly link it to a defined sequence on a bead at high efficiency using the same coupling procedure utilized in traditional CLIP-based approaches (see **Methods**). Using this pool, we perform on-bead immunopurification (IP) of RBPs in UV-crosslinked lysates using standard conditions and assign individual protein identities to their associated RNAs using split-and-pool barcoding, where the same barcode strings are added to both the oligonucleotide bead tag and immunopurified RNA (**Figure 1A**). We dramatically simplified our split-and-pool tagging method such that the entire protocol can be performed without the need for specialized equipment in ~1 hour (see **Methods**).

After split-and-pool tagging and subsequent library preparation, we sequenced all barcoded DNA molecules (antibody-bead tags and the converted cDNA of RNAs bound to corresponding RBPs). We then matched all antibody-bead tags and RNA reads by their shared barcodes; we refer to these as SPIDR clusters (**Figure 1A**). We merged all SPIDR clusters by protein identity (specified by the antibody-bead tag) to generate a high-depth binding map for each protein. The resulting datasets are analogous to those generated by traditional individual CLIP approaches.

To ensure that IP using a pool containing multiple antibodies can successfully and specifically purify each of the individual proteins, we performed an IP in K562 cells using a pool of antibodies against 39 RBPs and measured the purified proteins by liquid chromatography tandem mass spectrometry (LC-MS/ MS). We confirmed that 35 of the 39 targeted RBPs enriched at least 2-fold relative to a negative control, showing that multiplexed enrichment of several RBPs simultaneously is possible (**Supplemental Figure 2**). The few exceptions were RBPs that were simply not detected (neither in the pooled IP nor under control conditions) and likely reflect either a poor antibody or lack of RBP expression in this cell line.

### SPIDR accurately maps dozens of RBPs within a single experiment

To test whether SPIDR accurately maps RBPs to RNA, we performed SPIDR in two widely studied human cell lines (K562 and HEK293T cells). Specifically, we generated antibody bead pools containing 68 uniquely tagged antibody-beads targeting 62 distinct RBPs across the RNA life cycle, including splicing, processing, and translation factors (**Figure 1B, Supplemental Tables 1, 2**). As negative controls, we included antibodies against epitopes not present in endogenous human cells (GFP and V5), antibodies that lack affinity to any epitope (mouse IgG), and oligonucleotide-labeled beads lacking any antibody (empty beads).

Using these pools, we performed SPIDR on 10 million UV-crosslinked cells. Focusing on the K562 data (which were sequenced at greater depth), we generated a median of 4 oligonucleotide tags per SPIDR cluster with the majority of clusters (>80%) containing tags representing only a single antibody type (**Supplemental Figure 3**), indicating that there is minimal ‘crosstalk’ between beads in a SPIDR experiment. This specificity enables us to uniquely assign RNA molecules to their corresponding RBPs. After removing PCR duplicates, we assigned each sequenced RNA read to its associated RBP and identified high confidence binding sites by comparing read coverage across an RNA to the coverage in all other targets in the pooled IP (**Supplemental Figure 4, Supplemental Figure 5**; see **Methods** for details). Using this approach, we detected the precise binding sites for SAF-A, PTBP1, SPEN, and HNRNPK on the *XIST* RNA^17, 20, 23^ (**Figure 1C**). Although most proteins (38/53 RBPs in K562) contained more than 2 million mapped RNA reads (**Supplemental Figure 4**), we observed specific binding to known target sites even for RBPs with lower numbers of reads. For example, SLBP (Stem Loop Binding Protein) had only 1.5 million mapped reads yet displayed strong enrichment specifically at the 3’ ends of histone mRNAs as expected^29^ (**Figure 1D**).

To systematically assess the quality, accuracy, and resolution of our SPIDR binding maps and the scope of the SPIDR method, we explored several key features:

i. ***Accurate mapping of classical RNPs.*** We targeted RBPs of diverse functionality, such as those which bind preferentially to RNAs coding for proteins and/or lncRNAs, to introns, exons, miRNAs, etc., as well as more “classical” ribonuclear protein (RNP) complexes, such as the ribosome or spliceosome (**Figure 2A**). We observed precise binding to the expected RNAs and binding sites. For example, we observed binding of:

- LSM11 to the U7 small nuclear RNA (snRNA)^33^ and the telomerase RNA component (TERC)^34^ (**Figure 2A and 2B**).
- WDR43, a protein that is involved in ribosomal RNA (rRNA) processing, to the 45S pre-rRNA and the U3 small nucleolar RNA (snoRNA), which is involved in rRNA modification^35^ (**Figure 2A and 2C**).
- LIN28B to a distinct region of the 45S pre-rRNA, consistent with recent reports of its role in ribosomal RNA biogenesis in the nucleolus^36^ (**Figure 2A and 2C**).
- NOLC1 (also known as NOPP140), a protein that localizes within the nucleolus and Cajal bodies^37, 38^, to both the 45S pre-rRNA (enriched within the nucleolus) and various small Cajal-body associated RNAs (scaRNAs) (**Figure 2A**).
- DDX52, a DEAD-box protein that is predicted to be involved in the maturation of the small ribosomal subunit^39, 40^ and RPS3, a structural protein contained within the small ribosomal RNA subunit, to distinct sites on the 18S rRNA (**Figure 2A**).
- FUS and TAF15 to distinct locations on the U1 snRNA^41, 42^ (**Figure 2A**)
- SMNDC1 specifically to the U2 snRNA^43^ (**Figure 2A**)
- SSB (also known as La protein) binding to tRNA precursors consistent with its known role in the biogenesis of RNA Polymerase III transcripts^44, 45^ (**Figure 2A**).
- LIN28B to the let-7 miRNA^46–50^ (**Figure 2D)**,
- LARP7 binding to 7SK^51^ (**Figure 2A**, **Supplemental Figure 5**).
ii. ***Many RBPs bind their own mRNAs to autoregulate expression levels.*** Many RBPs have been reported to bind their own mRNAs to control their overall protein levels through post-transcriptional regulatory feedback^52–54^. For example, SPEN protein binds its own mRNA to suppress its transcription^55^, UPF1 binds its mRNA to target it for Nonsense Mediated Decay^56^, TARDBP binds its own 3’-UTR to trigger an alternative splicing event that results in degradation of its own mRNA^57, 58^, and DGCR8, which together with DROSHA forms the known microprocessor complex, binds a hairpin structure in *DGCR8* mRNA to induce cleavage and destabilization of the mRNA^59^ (**Figure 2E**). In addition to these cases, we observed autoregulatory binding of proteins to their own mRNAs for nearly a third of our targeted RBPs (15 proteins) (**Supplemental Figure 6**).
iii. ***Different antibodies that capture the same protein or multiple proteins within the same complex show similar binding***. We considered the possibility that including antibodies against multiple proteins contained within the same complex, or that otherwise bind to the same RNA, within the same pooled sample could compete against each other and therefore limit the utility of large-scale multiplexing. However, we did not observe this to be the case; in fact, antibodies against different proteins known to occupy the same complex displayed highly comparable binding sites on the same RNAs. For example, DROSHA and DGCR8, two proteins that bind as part of the microprocessor complex, showed highly consistent binding patterns across known miRNA precursors with significant overlap in their binding sites (odds-ratio of 316-fold, hypergeometric p-value < 10^−100^). Similarly, when we included two distinct antibodies targeting the same protein, HNRNPL, we observed highly comparable binding profiles for both antibodies (**Figure 2F**) and significant overlap in defined binding sites (odds-ratio of 15-fold, hypergeometric p-value < 10^−100^). Taken together, our results indicate that SPIDR can be used to map different RBPs that bind to the same RNA targets and can successfully map multiple antibodies targeting the same protein. As such, SPIDR may be a particularly useful tool for directly screening multiple antibodies targeting the same protein to evaluate utility for use in CLIP-like studies.
iv. ***Transcriptome-wide SPIDR maps are highly comparable with CLIP.*** Because K562 represents the ENCODE-mapped cell line with the largest number of eCLIP datasets, we were able to benchmark our SPIDR results directly to those generated by ENCODE. To do this, we compared the profiles for each of the 33 RBPs that overlap between SPIDR and ENCODE datasets in K562 cells^23, 28, 29^ (see **Methods**). We observed highly overlapping binding patterns for most RBPs, including: HNRNPK binding to *POLR2A* (**Figure 3A**), PTBP1 binding to *AGO1* (**Figure 3B**), RBFOX2 to *NDEL1* (**Figure 3C**) and the binding of several known nuclear RBPs to *XIST* (**Figure 3D**). To explore this data on a global scale, we compared RNA binding sites for each RBP and observed significant overlap between SPIDR- and ENCODE-derived binding sites for the vast majority of proteins (29/33, p<0.01, **Figure 3E**). Moreover, we observed that in virtually all cases each RBP preferentially binds to the same RNA features (e.g., introns, exons, CDS, miRNAs, 5’ and 3’UTRs) in both datasets (**Figure 3F**, **Supplemental Figure 7**). Finally, the binding motifs identified within the significant SPIDR-defined binding sites match those defined by CLIP and *in vitro* binding assays^29^ (e.g., RNA Bind-N-Seq, **Figure 3G**).
v. ***SPIDR enables high-resolution RBP mapping at single nucleotide resolution***. Next, we explored whether SPIDR can provide single nucleotide resolution maps of precise RBP-RNA binding sites, as is the case for some current CLIP-seq approaches. Specifically, UV crosslinking creates a covalent adduct at the site of RBP-RNA crosslinking, which leads to a preferential drop-off of the reverse transcriptase at these sites (**Figure 4A**). To explore this, we computed the number of reads that end at each position of an RNA (truncations) and compared these counts to those expected by chance. We observed strong positional enrichments at known protein binding sites. For example, we observe strong enrichment for RPS2 and RPS6 – two distinct structural components of the small ribosomal RNA subunit – at the precise locations where these proteins are known to contact the 18S rRNA in the resolved ribosome structure (**Figure 4B**). Moreover, examining individual mRNAs bound by HNRNPC (**Figure 4C**) or PTBP1 (**Figure 4D**) showed that the precise binding site corresponds to the known motif sequence. When we computed this enrichment more globally, we observed that HNRNPC (**Figure 4E**) and PTBP1 (**Figure 4F**) reads tend to terminate immediately proximal to these well-known binding sequences^29^.

**Figure 2:**
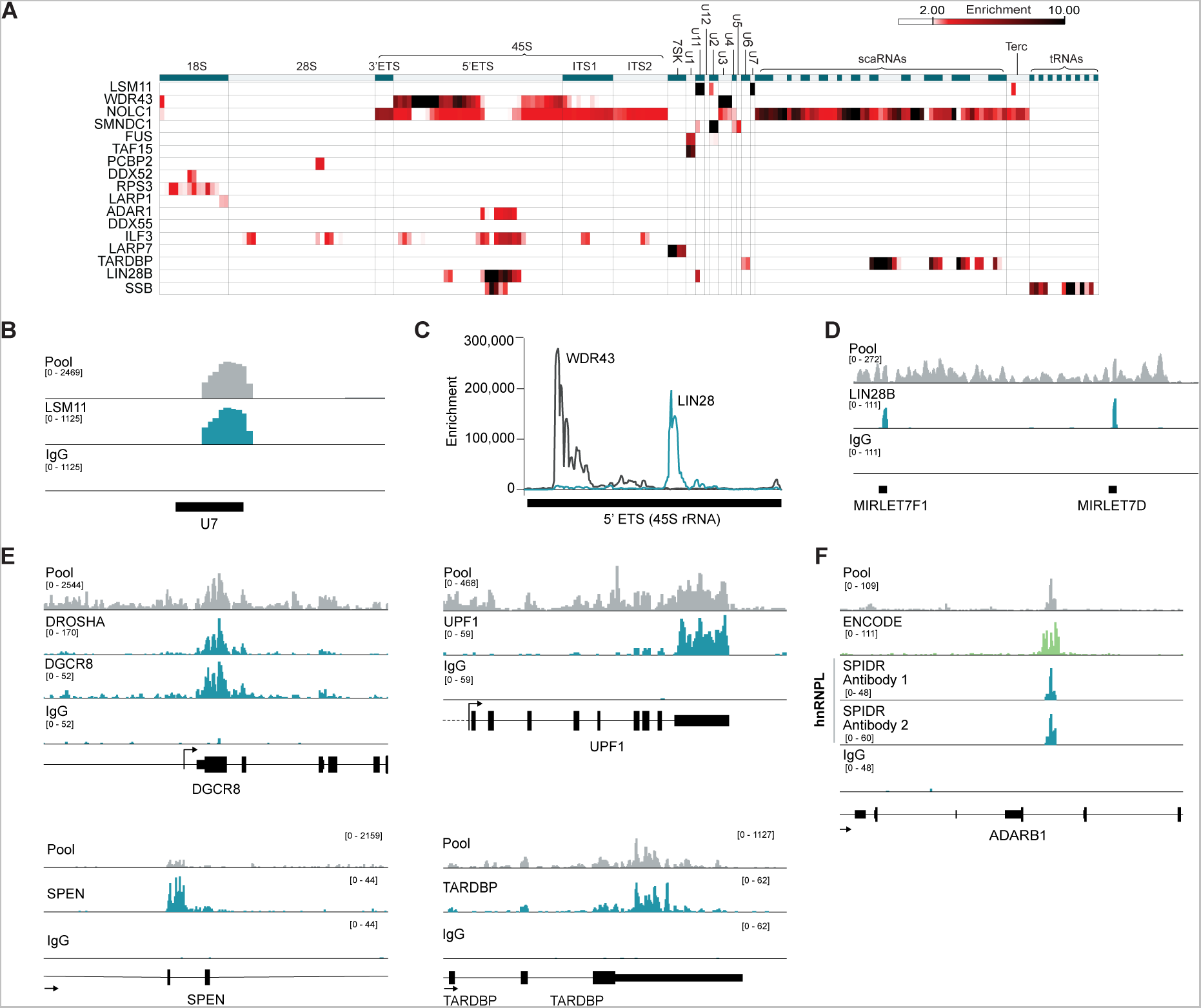
SPIDR accurately maps binding of a diverse set of RBPs. **(A)** RNA binding patterns of selected RBPs (rows) relative to 100nt windows across each classical non-coding RNA (columns). Each bin is colored based on the enrichment of read coverage per RBP relative to background. **(B)** Sequence read coverage for LSM11 binding to U7 snRNA. For all tracks, “pool” refers to all reads prior to splitting them by paired barcodes (shown in gray), and individual tracks (shown in teal) reflect reads after assignment to specific antibodies. **(C)** Enrichment of read coverage relative to background for WDR43 and LIN28B over the 5’ ETS region of 45S RNA. **(D**) Sequence reads coverage for LIN28B binding to let-7 miRNAs. **(E)** Sequence reads coverage for DROSHA/DGCR8, UPF1, SPEN, and TARDBP to their respective mRNAs. **(F)** Sequence reads coverage for two distinct antibodies to HNRNPL in a single SPIDR experiment. For comparison, HNRNPL coverage from the ENCODE-generated eCLIP data is shown (bright green).

**Figure 3:**
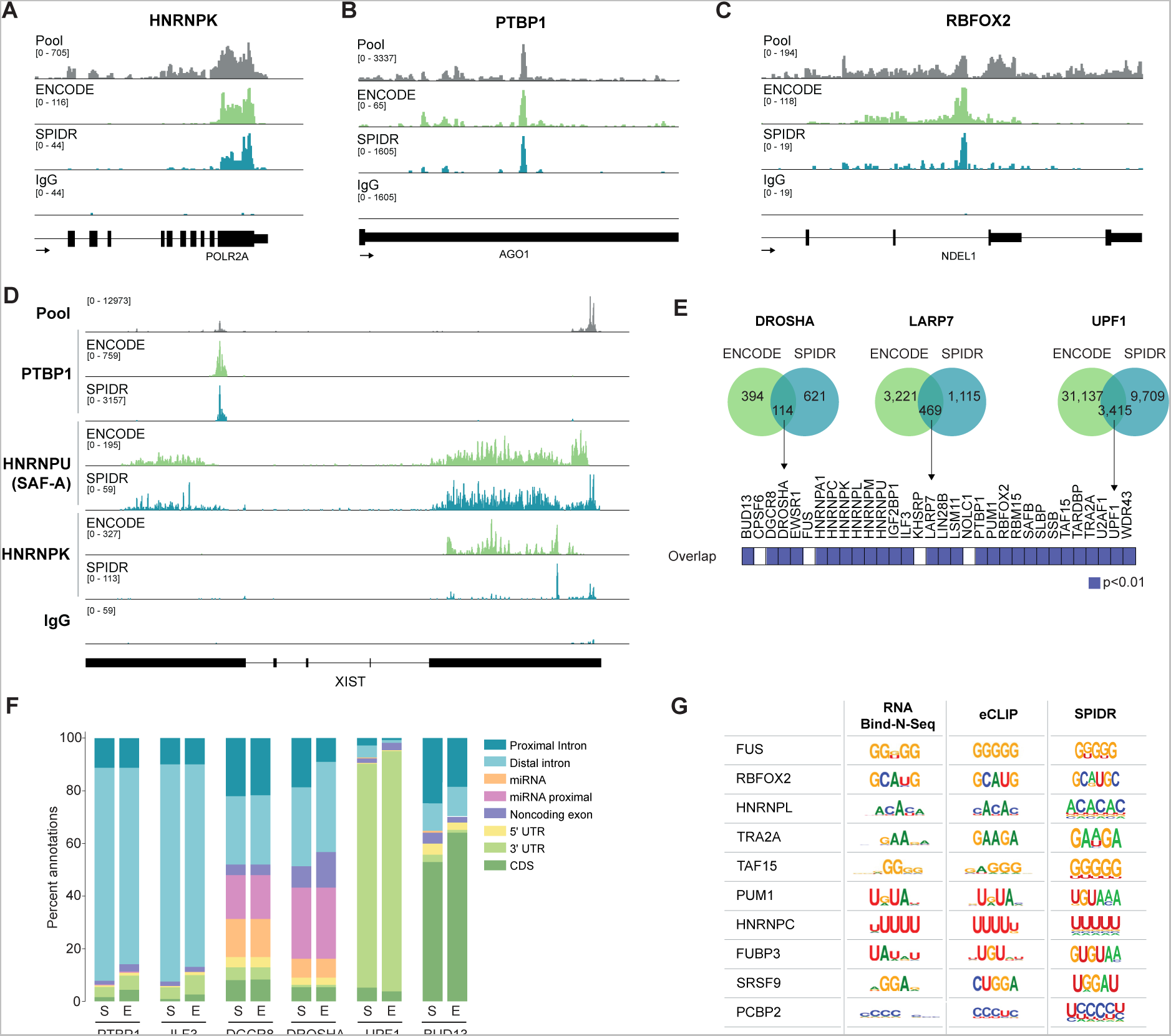
SPIDR data is highly comparable to previous eCLIP datasets. Examples of concordant binding identified by eCLIP (ENCODE consortium) and SPIDR. Sequence reads coverage is shown for individual proteins measured by ENCODE (green) and SPIDR (teal) along with a negative control (IgG). **(A)** HNRNPK, **(B**) PTBP1, and **(C)** RBFOX2. **(D**) Comparison of ENCODE and SPIDR data for multiple proteins bound to the *XIST* lncRNA. Sequence reads coverage for PTBP1, HNRNPU (SAF-A), and HNRNPK are shown. **(E)** Significance of overlap between binding sites detected by SPIDR and those identified within paired proteins in the ENCODE data. Each bin represents the paired protein between both experiments, blue represent a hypergeometric p-value of less than 0.01. **(F)** Peak annotation in matched SPIDR and ENCODE data. Stacked bar plot showing the percentage of peaks detected in the SPIDR (S) or ENCODE (E) datasets in various annotation categories. **(G)** Comparison of significant motifs identified within SPIDR peaks (right, p-value threshold < 1e-40) to those reported for RNA Bind-n-Seq (left) or eCLIP (middle)29.

**Figure 4:**
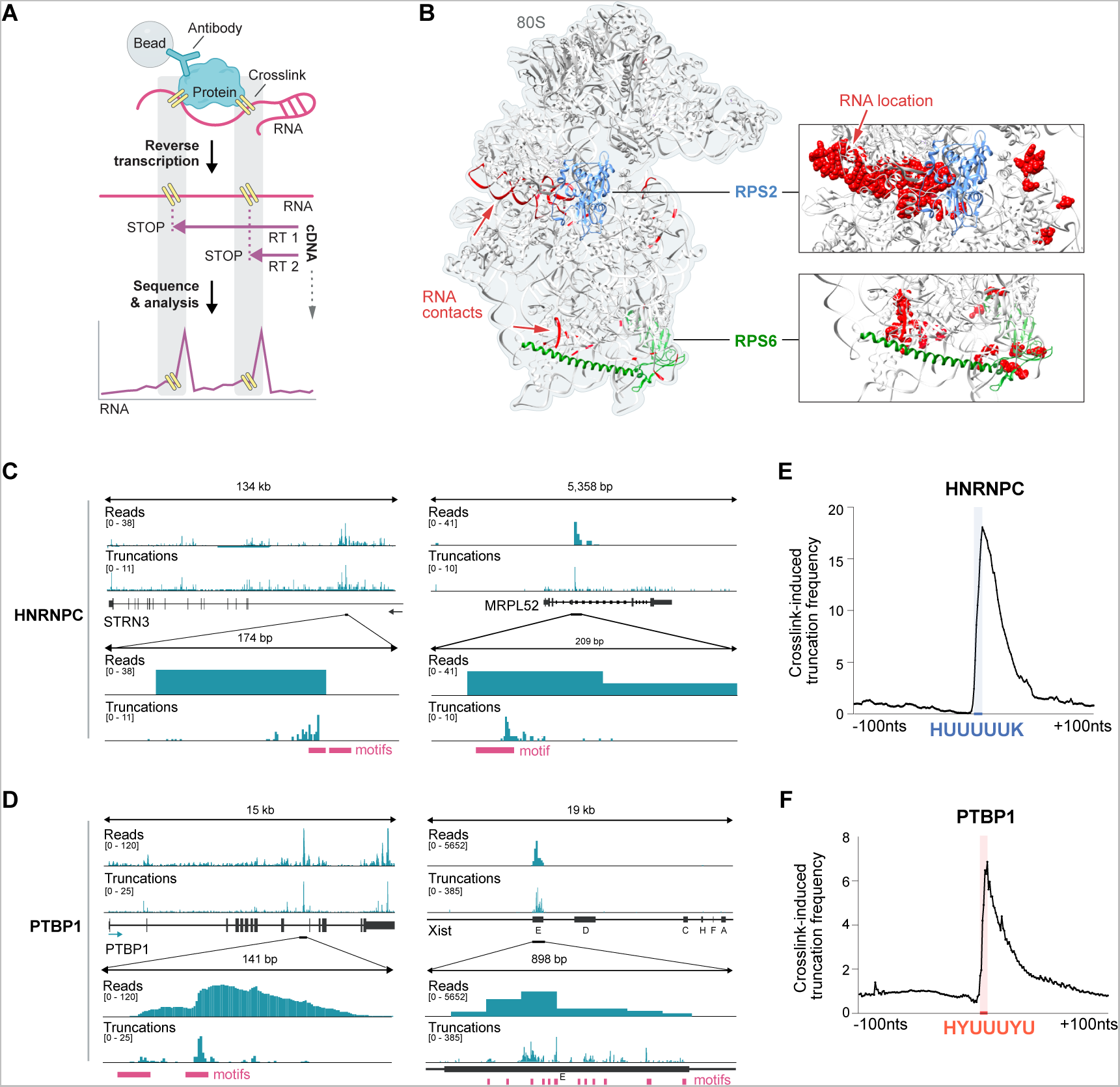
SPIDR enables high-resolution RBP mapping at single nucleotide resolution. **(A)** Schematic showing how reverse transcription pause sites can be used to map RBP-RNA interactions at single nucleotide resolution. UV light crosslinks the RBP to the target RNA at points of direct contact. During reverse transcription, the enzyme preferentially stalls at the crosslinking site, leading to termination of cDNA synthesis (STOP). Mapping the 3’-end of the cDNA (truncations) may identify the RBP binding site at single nucleotide resolution. **(B)** The SPIDR determined binding sites of RPS2 and RPS6 overlayed on the known 80S ribosome structure. RPS2 protein is shown in light blue and RPS6 protein in green; direct RNA contacts of each of these two ribosomal proteins detected by SPIDR are shown in red. SPIDR data shown is from HEK293T cells. **(C)** HNRNPC binding sites for *STRN3* (left) and *MRPL52* (right). Both raw read alignments (“Reads”, top) and 3’-end truncations of the cDNA (“Truncations”, bottom) are shown. The upper two panels show the mapped reads and truncations for the whole gene, the lower two panels are zoomed-in on the indicated region. The known binding motifs for HNRNPC are depicted in magenta. **(D**) Examples of PTBP1 binding sites for *PTBP1* itself (left) and *XIST* (right). Both raw read alignments and 3’-end truncations of the cDNA reads are shown. Known binding motifs for PTBP1 are in magenta. **(E)** Truncation frequency (3’ ends of the mapped cDNA reads) over all significantly enriched HNRNPC peaks is shown centered on the motif position. The region of the steep frequency rise of truncations is shown by the blue line on the y-axis and corresponds to the sequence shown in blue. **(F)** The truncation frequency (3’ ends of the mapped cDNA reads) over all significantly enriched PTBP1 peaks is shown relative to the motif position within each peak. The region of the steep frequency rise of truncations is shown by the orange line on the y-axis and corresponds to the sequence shown in orange.

Taken together, our data demonstrate that SPIDR generates highly accurate single nucleotide RBP-binding maps for dozens of RBPs within a single experiment. Moreover, SPIDR can simultaneously map RBPs representing diverse functions and binding modalities, including RBPs that bind within thousands of RNAs (e.g. CPSF6), RBPs that bind only a few very specific RNAs (e.g. SLBP), as well as RBPs that bind primarily within intronic regions within the nucleus (e.g. PTBP1) and RBPs that bind primarily to exonic regions within the cytoplasm (e.g. UPF1).

### LARP1 binds to the 40S ribosome and mRNAs encoding translation-associated proteins

In addition to the three known structural components of the small ribosomal subunit (RPS2, RPS3, and RPS6), we noticed that LARP1 also showed strong binding to the 18S ribosomal RNA (**Figures 2A, 5A**). LARP1 is an RNA binding protein that has been linked to translational initiation of specific mRNAs. It is known to bind to the 5’ end of specific mRNAs, primarily those encoding critical translation proteins such as ribosomal proteins and initiation and elongation factors, via recognition of a terminal oligopyrimidine (TOP) sequence in the 5’ UTR of these transcripts^60^. The exact role of LARP1 in translation has been debated because it has been reported to both promote and repress translation of mRNAs containing a TOP-motif^60–66^.

**Figure 5:**
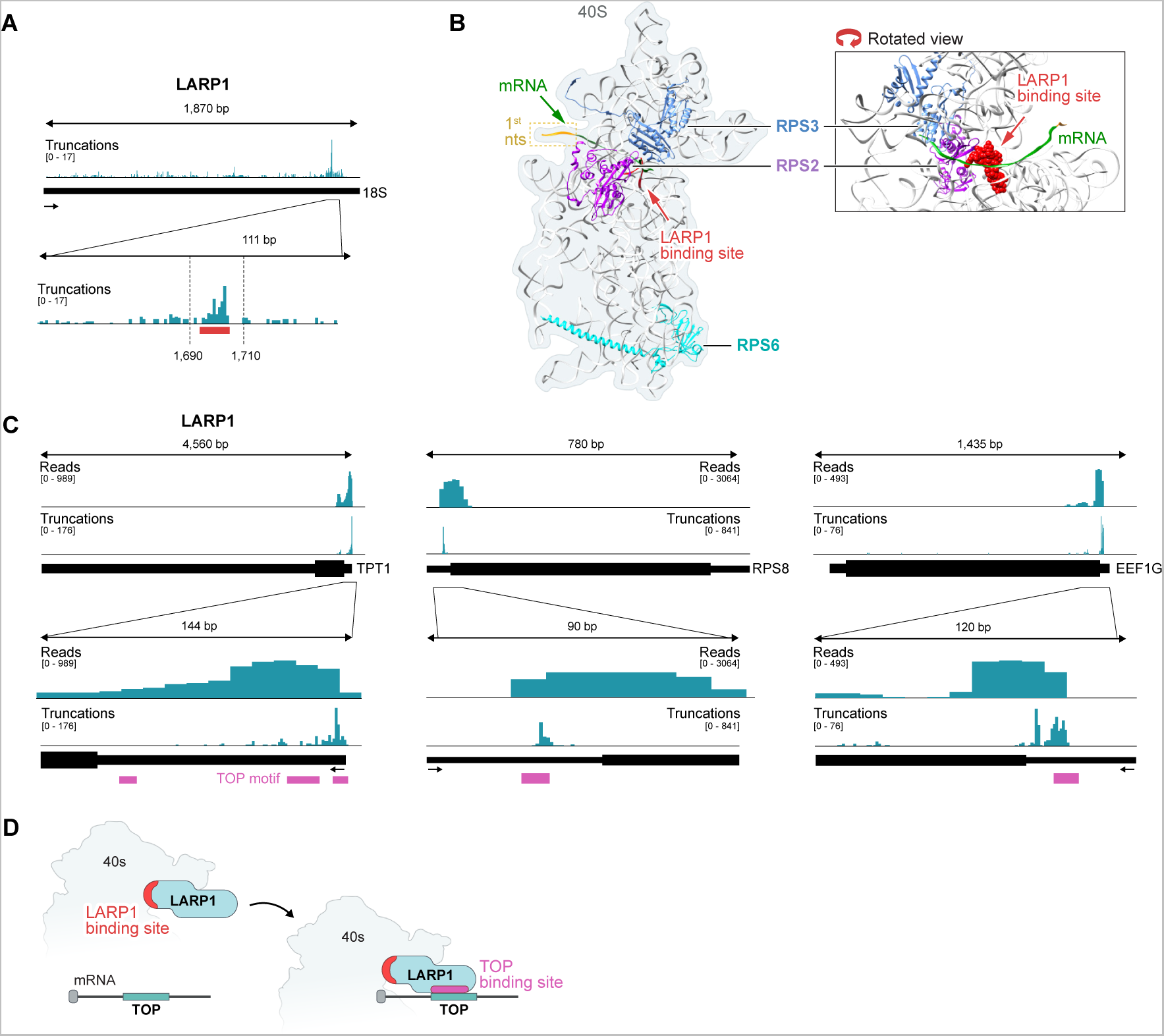
LARP1 binds to 18S rRNA near the mRNA entry channel and at TOP-motifs contained within the 5’UTRs of mRNAs. **(A)** Frequency of 3’-end truncations of LARP1 reads plotted across the 18S rRNA. Zoom-in shows accumulation near nucleotide position 1700 (indicated by the red bar). Data shown is from HEK293T cells. **(B)** The structure of the 40S ribosomal subunit bound to the 5’-end of an mRNA molecule (green). The first nucleotide of the mRNA is indicated in orange. RPS2 (purple) and RPS3 (blue) are indicated for orientation and the mRNA is shown in green. The LARP1 binding site detected on the 18S rRNA is indicated in red. **(C)** Examples of LARP1 binding for three different mRNAs containing TOP motifs in their 5’UTRs – *TPT1* (left), *RPS8* (middle) and *EEF1G* (right). Both read alignments (“Reads”, top) and 3’end truncations of the cDNA reads (“Truncations”, bottom) are shown. TOP motifs within the 5’-UTRs are depicted in magenta. **(D)** Model of LARP1 interactions based on SPIDR data. LARP1 shows preferential binding to both the 18S rRNA (close to the mRNA entry channel of the 40S subunit) and the TOP motifs within the 5’UTR of specific mRNAs. In this way, LARP1 could facilitate recruitment of the 40S subunit to TOP motif-containing mRNAs.

Although LARP1 is known to bind TOP-motif containing mRNAs, how it might promote translation initiation of these mRNAs is mostly unknown. Because we identified a strong binding interaction between LARP1 and the 18S ribosomal RNA, we explored where in the initiating ribosome this interaction occurs. Interestingly, the LARP1 binding site on the 18S ribosomal RNA (1698-1702 nts) is at a distinct location relative to all other 18S binding proteins that we explored and corresponds to a position within the 48S structure that is directly adjacent to the mRNA entry channel (**Figure 5B**). More generally, we observed strong binding of LARP1 at the TOP-motif sequence within the 5’ UTR of translation-associated mRNAs (**Figure 5C**).

These results suggest that LARP1 may act to promote increased translational initiation of TOP-motif containing mRNAs by directly binding to the 43S pre-initiation complex and recruiting this complex specifically to mRNAs containing a TOP-motif. Because LARP1 is positioned immediately adjacent to the mRNA in this structure, this 43S+LARP1 complex would be ideally positioned to access and bind the TOP motif to facilitate efficient ribosome assembly and translational initiation at these mRNAs. This mechanism of direct ribosome recruitment to TOP-motif containing mRNAs through LARP1 binding to the 43S ribosome and the mRNA would explain why the TOP-motif must be contained within a fixed distance from the 5’ cap to promote translational initiation^67^ (**Figure 5D**).

### 4EBP1 binds specifically to LARP1-bound mRNAs upon mTOR inhibition

Translation of TOP motif-containing mRNAs is selectively repressed upon inhibition of the mTOR kinase, which occurs in conditions of physiological stress^68–71^. Recent studies have shown that under these conditions, LARP1 binds the 5’-UTR of TOP-containing mRNAs, and it has been postulated that this binding activity is responsible for the specific translational repression of these mRNAs^60, 72^. Yet, the mechanism by which LARP1 binding might repress translation remains unknown.

The canonical model for how mTOR inhibition leads to translational suppression is through the selective phosphorylation of 4EBP1^69, 71^. Specifically, when phosphorylated, 4EBP1 cannot bind to EIF4E, which is the critical initiation factor that binds to the 5’ mRNA cap and recruits the remaining initiation factors through direct binding with EIF4G^73, 74^. When 4EBP1 is not phosphorylated (i.e., in the absence of mTOR), it binds to EIF4E and prevents it from binding to EIF4G and initiating translation. While this differential binding of 4EBP1 to EIF4E upon mTOR modulation is well-established and is central to translational suppression, precisely how it leads to selective modulation of TOP mRNA translation has remained unclear. Specifically, direct competition between 4EBP1 and EIF4G for binding to EIF4E should impact translation of all EIF4E-dependent mRNAs, yet the observed translational downregulation is specific to TOP-containing mRNAs^69, 71, 75^ and this specificity is dependent on LARP1 binding^60^.

To explore the mechanism of translational suppression of TOP-containing mRNAs upon mTOR inhibition, we treated HEK293T cells with torin, a drug that inhibits mTOR kinase. We adapted SPIDR to map multiple independent samples within a single split-and-pool barcoding experiment (**Figure 6A, Supplemental Table 2,** see **Methods**) and used this approach to perform SPIDR on >50 distinct RBPs, including LARP1, numerous translational initiation factors, and 4 negative controls in both torin-treated and untreated conditions.

**Figure 6:**
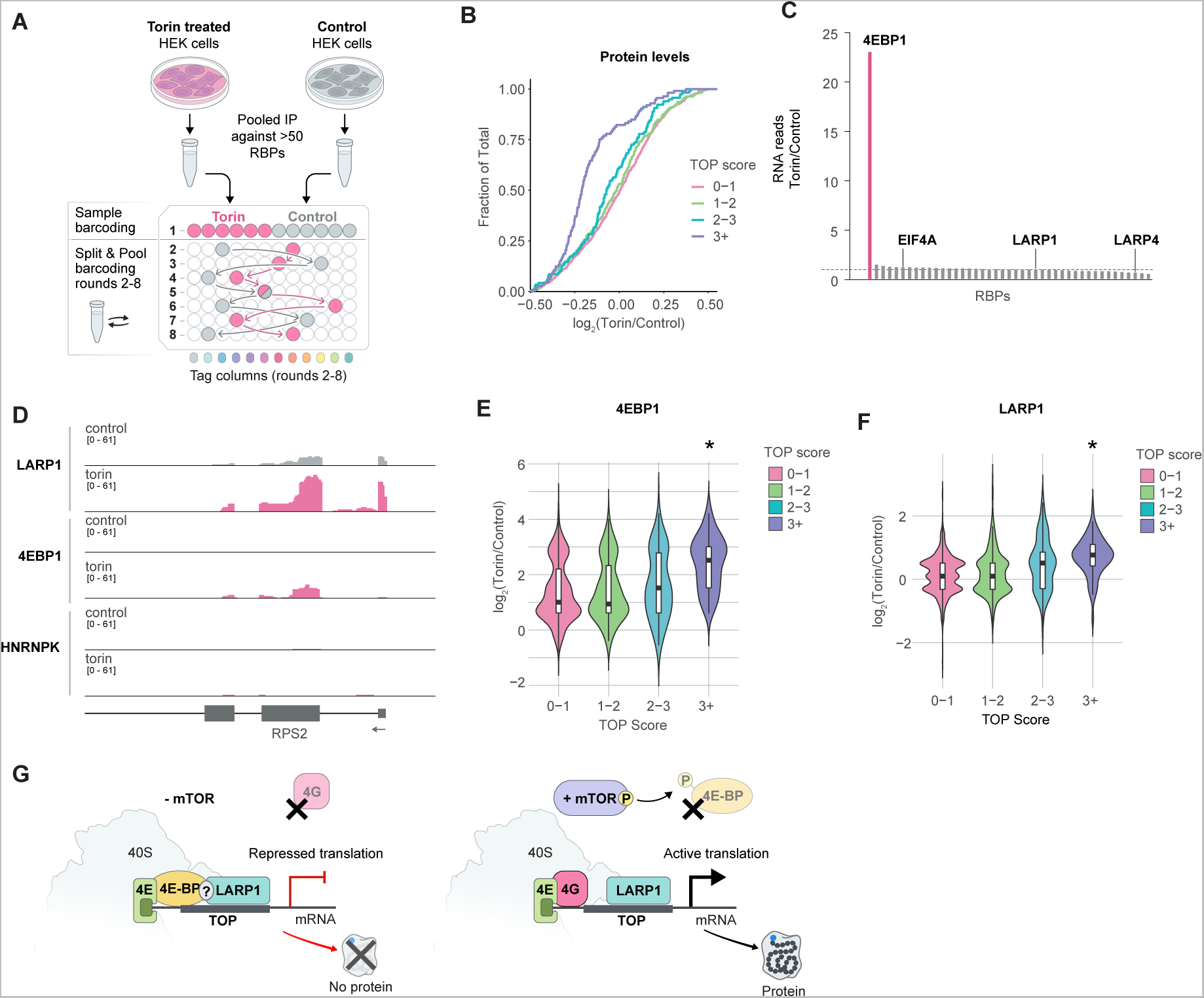
4EBP1 binds specifically to LARP1-bound mRNAs upon mTOR inhibition. **(A)** Schematic of experimental approach for the mTOR perturbation experiment. HEK293T cells were treated with either 250nM torin or control (solvent only) for 18 hours. SPIDR was performed on both samples. The multiplexed IP was performed separately, and the samples were mixed after the first round of barcoding (see Methods for details). **(B)** Cumulative Distribution Function (CDF) plots of protein changes in torin versus control treated samples as determined by LC-MS/MS. log2 ratios (Torin/Control) are shown on the x-axis and fraction of total (from 0 to 1) is shown on the y-axis. Proteins were grouped into four categories based on their TOP motif score as previously published in (Philippe et al., 2020)60. The analysis was performed on the 2000 most highly expressed genes (based on RNA expression, see Methods). **(C)** Number of SPIDR reads assigned to each RBP in the torin-treated samples versus control samples. 4EBP1 (pink line), EIF4A, LARP1 and LARP4 are also indicated. Dashed line corresponds to enrichment of 1. **(D)** Raw alignment data for selected RBPs across *RPS2*, an mRNA with a strong TOP motif. For each protein “control” and “torin” treatment tracks are shown. **(E)** Violin plots of the log2 ratios (torin/control) of significant binding sites for 4EBP1 are shown. The RNA targets are grouped based on their TOP motif score as published in (Philippe et al., 2020)60. **(F)** Violin plots of the log2 ratios (Torin/Control) of significant binding sites for LARP1 are shown. The RNA targets were grouped based on their TOP motif score as published in (Philippe et al., 2020)60. For (E) and (F) the asterisks indicate statistical significance (p-value < 0.00001, Mann-Whitney). **(G)** Model of mTOR-dependent repression of mRNA translation. LARP1 binds to the 40S ribosome and to 5’ untranslated region of TOP-containing mRNAs independent of mTOR activity. When mTOR is active (i.e., in the absence of torin; right side), this dual binding modality can recruit the ribosome specifically to TOP-containing mRNAs and promote their translation. When mTOR is inactive (i.e., in the presence of torin), 4EBP1 can bind to TOP-containing mRNAs (possibly through an interaction with LARP1) and to EIF4E. The interaction between 4EBP1 and EIF4E prevents binding between EIF4E and EIF4G, which is required to initiate translation. In this way, LARP1/4EBP1 binding specifically to TOP-containing mRNAs would enable sequence-specific repression of translation.

To ensure that mTOR inhibition robustly leads to translational suppression of TOP-containing mRNAs, we quantified global protein levels in torin-treated and untreated cells using quantitative mass spectrometry (see **Methods**) to determine protein level changes globally. Although the level of most proteins does not change upon torin-treatment, we observed a striking reduction of proteins encoded from TOP motif-containing mRNAs. Indeed, this translational suppression was directly proportional to the strength of the TOP-motif contained within the 5’-UTR of each mRNA (**Figure 6B**).

Next, we explored changes in RBP binding upon mTOR inhibition. We measured the number of RNA reads observed for each protein upon torin treatment relative to control. While the majority of proteins showed no change in the number of RNA reads, the sole exception was 4EBP1, which showed a dramatic increase (>20-fold) in the overall number of RNA reads produced upon mTOR inhibition (**Figure 6C**). Interestingly, this increase corresponded to increased binding specifically at mRNAs containing a TOP-motif (p-value < 8 x 10^−10^, Mann-Whitney, **Figure 6D** and **6E**). Notably, this did not simply reflect an increased level of 4EBP1 binding at the same sites, but instead corresponded to the detection of many statistically significant binding sites only upon mTOR inhibition that were not observed in the presence of mTOR activity (control samples). Consistent with these observations, a previous study observed that 4EBP1 can be in proximity to translationally suppressed mRNAs upon mTOR inhibition^76^.

In contrast to 4EBP1, which showed a dramatic transition in binding activity to mRNA upon mTOR inhibition, we did not observe a global change in the number of RNA reads purified by LARP1 upon mTOR inhibition (**Figure 6C**). Indeed, in both torin-treated and untreated samples we observed strong binding of LARP1 to TOP motif mRNAs as well as to the 18S ribosomal RNA suggesting that this interaction with the 40S ribosome and TOP mRNAs occurs independently of mTOR activity. However, we did observe a 1.7-fold increase in levels of binding of LARP1 at TOP mRNAs upon mTOR inhibition (p-value < 5.4 x 10^−16^, Mann-Whitney, **Figure 6D** and **6F**). This increased enrichment at TOP mRNAs could reflect more LARP1 binding at these specific mRNAs or could reflect the fact that the LARP1 complex might be more stably associated with each mRNA due to translational repression.

Together, our results suggest a model that may reconcile the apparently divergent perspectives about the role of LARP1 as both an activator and repressor of translational initiation and explains how selective mTOR-dependent translational repression is achieved (**Figure 6G**). Specifically, LARP1 binds to the 40S ribosome and 5’ untranslated region of mRNAs containing a TOP motif regardless of mTOR activity. In the presence of mTOR (**Fig 6G, right side**), this dual binding modality can act to promote ribosome recruitment specifically to TOP-containing mRNAs and promote translation of these mRNAs. In the absence of mTOR (**Fig 6G, left side**), 4EBP1 can bind to TOP-containing mRNAs, potentially via the LARP1 protein already bound to these mRNAs. Indeed, most of the significant 4EBP1 binding sites are also bound by LARP1 under Torin treatment (60% overlap, odds-ratio of 12-fold, hypergeometric p-value < 10^−100^). By binding selectively to these TOP-containing mRNAs, 4EBP1 can bind to EIF4E and prevent binding between EIF4E and EIF4G, a necessary requirement for initiation of translation. In this way, LARP1/4EBP1 binding to specific mRNAs would enable sequence-specific repression of mRNA translation. This model would explain the apparently divergent roles of LARP1 as both an activator and repressor of translation as it indicates that LARP1 may act as a selective recruitment platform that can either activate or repress translation through the distinct factors that co-bind in the presence or absence of mTOR activity.

## DISCUSSION

Here we present SPIDR, a massively multiplexed method to generate high-quality, high-resolution, transcriptome-wide maps of RBP-RNA interactions. SPIDR can map RBPs with a wide-range of RNA binding characteristics and functions (e.g., mRNAs, lncRNAs, rRNAs, small RNAs, etc.) and will enable the study of diverse RNA processes (e.g., splicing, translation, miRNA processing, etc.) within a single experiment and at an unprecedented scale.

While we show that SPIDR can accurately map dozens of RBPs within a single experiment, the numbers used mostly reflect the availability of high-quality antibodies. As such, we expect that this approach can readily be applied to even larger pool sizes for hundreds or thousands of proteins simultaneously. Because of this, we expect that SPIDR will represent a critical technology for exploring the many thousands of human proteins that have been reported as putative RNA binding proteins but that remain largely uncharacterized^6–10^. Similarly, we expect that this technology will be crucial for assessing the putative functions of the >20,000 annotated ncRNAs which have remained largely uncharacterized.

Because the number of cells required to perform SPIDR is comparable to that of a traditional CLIP experiment, yet a single SPIDR experiment reports on the binding behavior of dozens (and likely hundreds) of RBPs, this approach dramatically reduces the number of cells required to map an individual RBP. Accordingly, SPIDR will be a valuable tool for studying RBP-RNA interactions in many different contexts, including within rare cell types and patient samples where large numbers of cells may be difficult to obtain.

We showed that SPIDR generates single nucleotide contact maps that accurately recapitulate the RNA-protein contacts observed within structural models. This suggests that SPIDR will also be well-suited to add high-resolution binding information for entire RNP complexes in a single experiment, as it will allow simultaneous targeting of all proteins within a complex. We envision that, in conjunction with more traditional structural biology methods, this approach will help elucidate the precise structure of various RNP complexes, including for mapping proteins that are not currently resolved within these structures (e.g. LARP1 binding within the 48S ribosome).

In addition to accurately measuring multiple proteins simultaneously, because of the nature of the split-and-pool barcoding strategy used, this approach also allows for multiple samples to be pooled within a single experiment. This ability to simultaneously map multiple proteins across different samples and conditions will enable exploration of RBP binding patterns and their changes across diverse biological processes and disease states. Until now, systematic comparative studies of RBP-RNA interaction changes at scale have been impossible, even for large consortia (e.g., ENCODE), which have invested massive amounts of time and effort to generate CLIP-seq data for only two cell lines. Our 4EBP1 results highlight the critical value of SPIDR for enabling exploration of RBP dynamics across samples. Specifically, 4EBP1 was not commonly thought to directly bind to mRNA, nonetheless, including 4EBP1 within our larger pool of target proteins allowed us to uncover changes across two different experimental conditions that may explain how specificity of mTOR-mediated translational suppression is achieved.

Although we focused on the differential RNA binding properties of 4EBP1/LARP1, there are many additional insights into RBP biology that we expect can be uncovered from exploration of this dataset. For example, we observe that TARDBP (TDP43) shows strong binding to U6 snRNA and to multiple scaRNAs, a class of ncRNAs that play critical roles in spliceosome-associated snRNA biogenesis^77^. TDP43 is an RBP of great interest because of its well-known genetic link to various neurodegenerative disorders, such as amyotrophic lateral sclerosis (ALS)^78–81^. These observations could provide new mechanistic insights into how disruption of this RBP impacts splicing changes and pathogenesis in neurodegeneration.

Thus, we expect that SPIDR will enable a fundamental shift for studying mechanisms of transcriptional and post-transcriptional regulation. Rather than depending on large consortium efforts to generate reference maps within selected cell-types, SPIDR enables any standard molecular biology lab to rapidly generate a comprehensive and high-resolution genome-wide map within any cell-type or experimental system of interest without the need for specialized training or equipment.

## ACKNOWLEDGEMENTS

We thank all members of the Jovanovic and Guttman labs for technical help as well as helpful comments about the manuscript, Shawna Hiley for editing, and Inna-Marie Strazhnik for illustrations. This work was funded by grants from NIH (R35GM128802 (MJ); R01AG071869 (MJ) R01HG012216 (MJ/MG), U01DK127420 (MG), R01DA053178 (MG), and F30CA278005 (JKG)), NSF (Award 2224211 (MJ)), CZI Ben Barres Early Career Acceleration Award (MG), and funds from Columbia University (MJ).

## NOTES

### Note 1: Comparison to a previous multiplexed CLIP method

A recent study reported a variant of CLIP called Antibody-Bead eCLIP (ABC) that utilizes direct chemical conjugation of an oligo sequence to an antibody followed by proximity ligation between the antibody-oligo and RNA to enable multiplexed mapping of 10 proteins simultaneously^27^. Our approach differs from this strategy in several key practical and conceptual ways.

***Antibody labeling*:** The ABC method utilizes direct chemical modification of each antibody. First, this requires large excess of each antibody and selective purification of each conjugate to generate each labeled reagent. This necessitates a more elaborate, multi-step chemical modification and purification procedure for each antibody and therefore is not readily accessible for labeling large numbers of distinct antibodies. Second, because ABC utilizes chemical modification of the antibody using NHS chemistry, the precise site of oligo conjugation on each antibody is random. This could impact both epitope recognition (when modified within the recognition site) and protein-G binding to the FC region of the antibody; both will decrease the efficiency of IP. In contrast, SPIDR utilizes labeling of the protein G bead instead of direct modification of the antibody. As such, SPIDR is a rapid, efficient, and highly modular strategy already utilized in standard IP strategies to couple antibodies to beads. Because of this distinction, the SPIDR approach can work with the same amounts of antibody used in standard approaches and with antibodies produced and stored in any buffer condition, without the need for specific purification or chemical modification.

***Proximity-ligation versus split-pool detection*:** The ABC method utilizes proximity-ligation to link an antibody sequence to its RNA target. There are several conceptual limitations to this strategy. First, the efficiency of proximity-ligation is limited as ligation must occur between each oligo and RNA end at 1:1 stoichiometry, resulting in many failed ligation events. This will decrease the efficiency of the overall RNA detection rate, an issue that will primarily impact RNAs of low abundance or in low cell numbers. Second, proximity-ligation methods are highly sensitive to distance constraints between the two ligating components. Accordingly, the success of this approach will depend on the distance between the RBP and the RNA and where on the RBP the specific antibody binds. Therefore, there are likely to be antibodies for which this approach will not produce comparable results to standard CLIP. Moreover, this approach is highly sensitive to the RNA fragment size generated. If fragments are too long, this will be problematic because there might be multiple proteins that can ligate; if the fragments are too short, the ends might not be capable of ligation. In contrast, the SPIDR method utilizes split-pool barcoding which is not dependent on the distances between the antibody, RBP, and RNA and therefore not susceptible to these distance constraints. Because of this, SPIDR could be used for analyzing RBP-RNA interactions within higher-order assemblies and in the presence of additional crosslinkers beyond UV. Finally, because SPIDR utilizes barcodes on the beads and because split-pool barcoding is not limited to pairwise contacts, a single barcode can provide information on the identity of multiple RNAs simultaneously thereby increasing the resolution detected per sequenced read.

For these reasons, we believe that SPIDR will be more broadly applicable to wide-range of antibodies and is readily accessible to any molecular biology lab.

### Note 2: Features and Limitations of the Method

First, similar to CLIP and other immunoprecipitation methods, SPIDR is constrained by the availability of antibodies that have been validated to specifically enrich for RBPs of interest. We note that SPIDR may offer the opportunity to partly alleviate this problem as its multiplexing capability allows for the inclusion of several distinct antibodies, including those that may not have been previously validated, against an RBP of interest without increasing the experimental burden.

Second, the SPIDR protocol requires that each experiment is performed under the same IP conditions for all RBPs. Although we show that standard conditions work for many diverse proteins, they may not be suitable for all RBPs. One possible solution is to match antibodies (and target RBPs) by similar IP conditions.

Finally, in the current protocol, we used the same antibody amount for each RBP of interest, which may in part explain the uneven coverage of RNA reads measured for each RBP. Although we do identify well-known binding sites for nearly all RBPs targeted, higher sequencing coverage might be needed for RBPs at the lower end of this distribution. An alternative solution would be to adapt the antibody amount to equalize coverage for each RBP after performing an initial pre-screen and low-depth sequencing run.

---

**Supplemental Figure 1:**
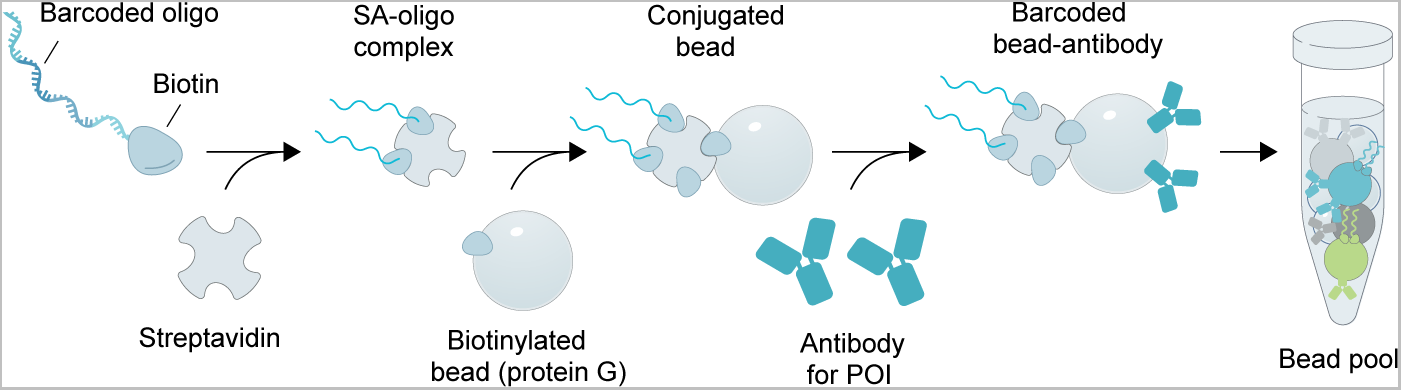
Schematic of our multiplexed antibody-bead labeling strategy. Populations of biotinylated protein G beads are incubated with a streptavidin-biotin oligo complex. Each population of beads is labeled with an oligo with a specific sequence and then incubated with one type of capture antibody such that each population has a unique capture antibody and a corresponding oligo tag that can be recognized after sequencing. Populations are combined to create the bead pool.

**Supplemental Figure 2:**
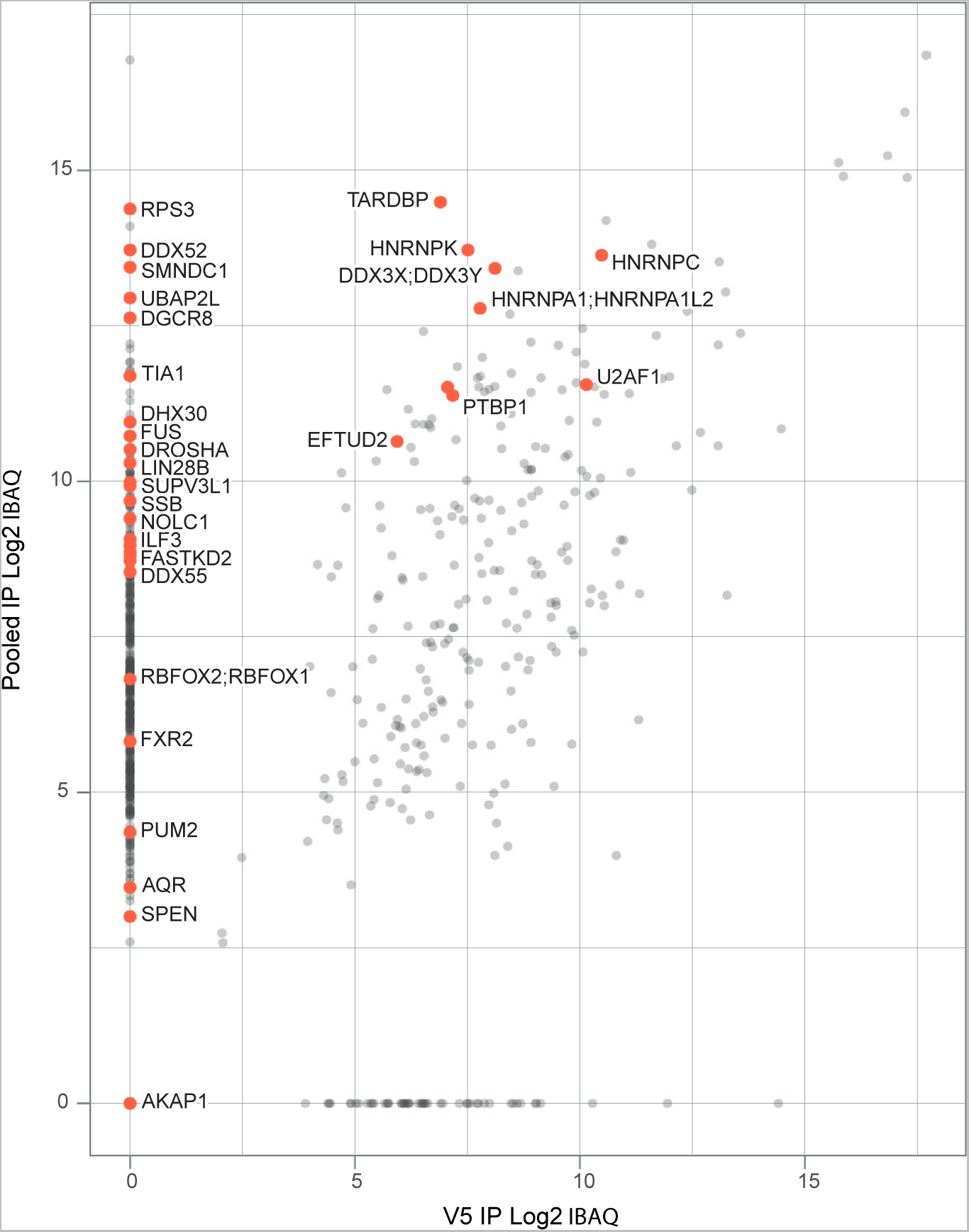
Multiplexed IP of dozens of RBPs accurately recovers targeted proteins. Scatter plot showing log2 transformed IBAQ (intensity based absolute quantification)82 values for all identified proteins in either the pooled IP with 39 targets (y-axis) versus those detected with a V5 negative control IP (x-axis) by LC-MS/MS. Target proteins that should be detected by the antibodies included in the pool of 39 used are marked in red.

**Supplemental Figure 3:**
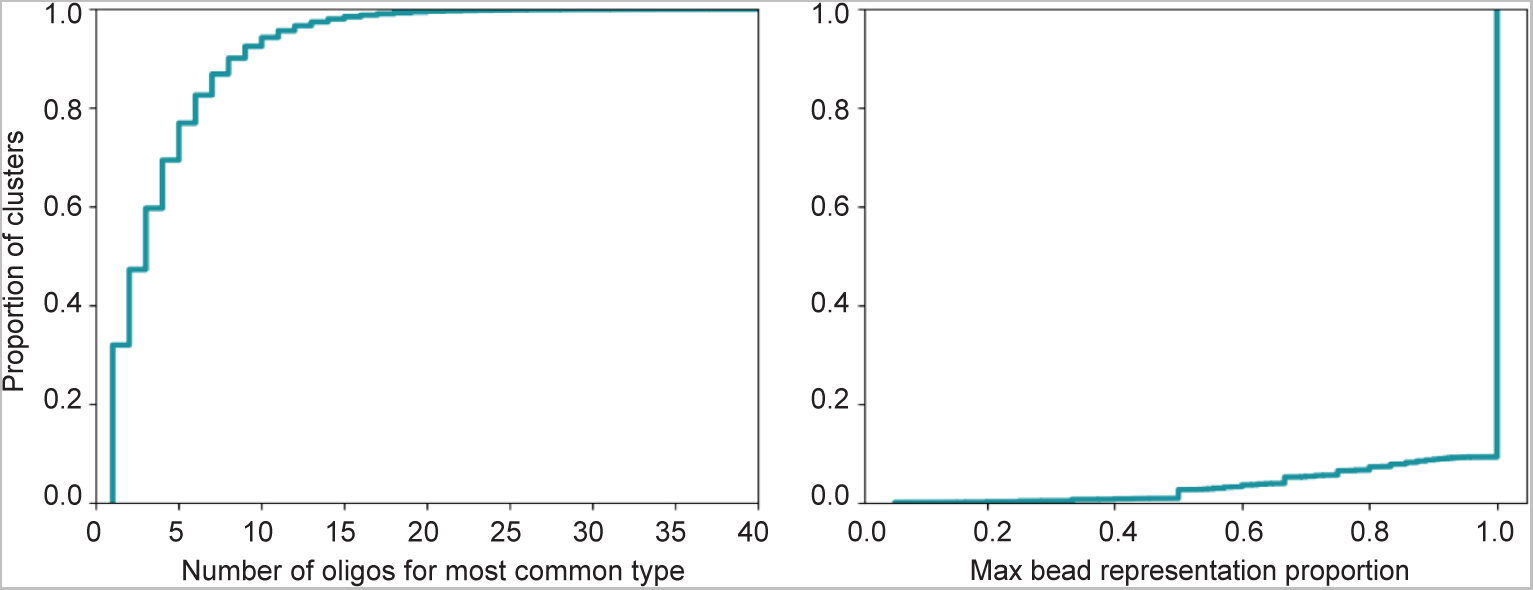
Uniqueness of beads and number of oligos per bead of the experiment. Observed distributions of labeled beads after sequencing. Each bead is defined in sequencing by a particular, unique combinatorial barcode acquired during split-pool. A SPIDR cluster represents any set of molecules, oligo or RNA, that share the same bead combinatorial barcode. Left: CDF plot showing the number of independent oligos matched within an individual SPIDR cluster. Right: CDF plot describing the degree of heterogeneity of these detected oligos within each SPIDR cluster, as determined by oligos with a shared combinatorial barcode. X axis represents the homogeneity of the oligo types with 1 indicating that all oligos are of the same type.

**Supplemental Figure 4:**
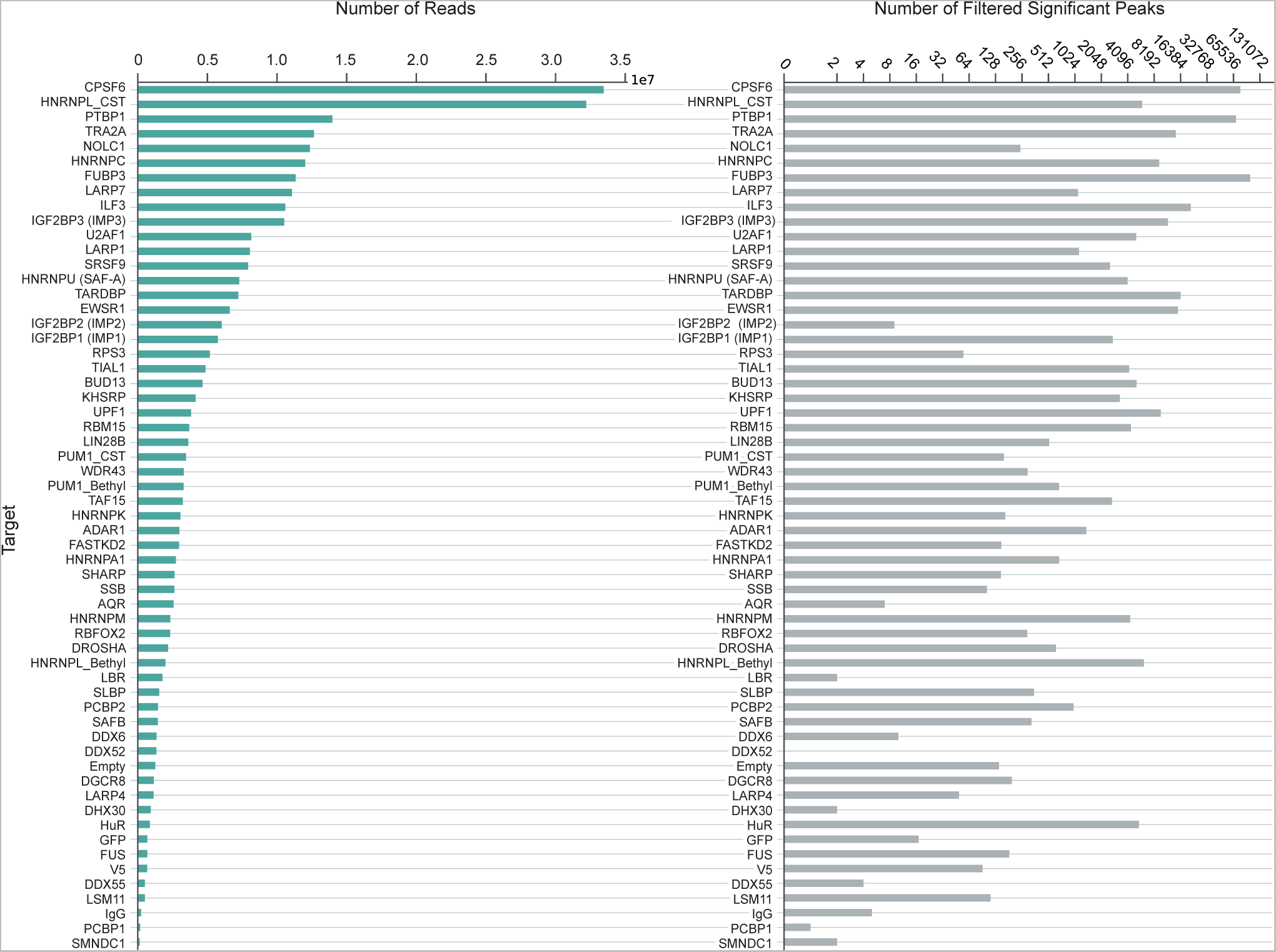
Mapped unique reads per RBP and significant binding sites identified per RBP. Number of deduplicated mapped reads and number of significant binding sites within uniquely mapped genomic regions per IP. The order is determined by the number of unique mapped reads in both plots.

**Supplemental Figure 5:**
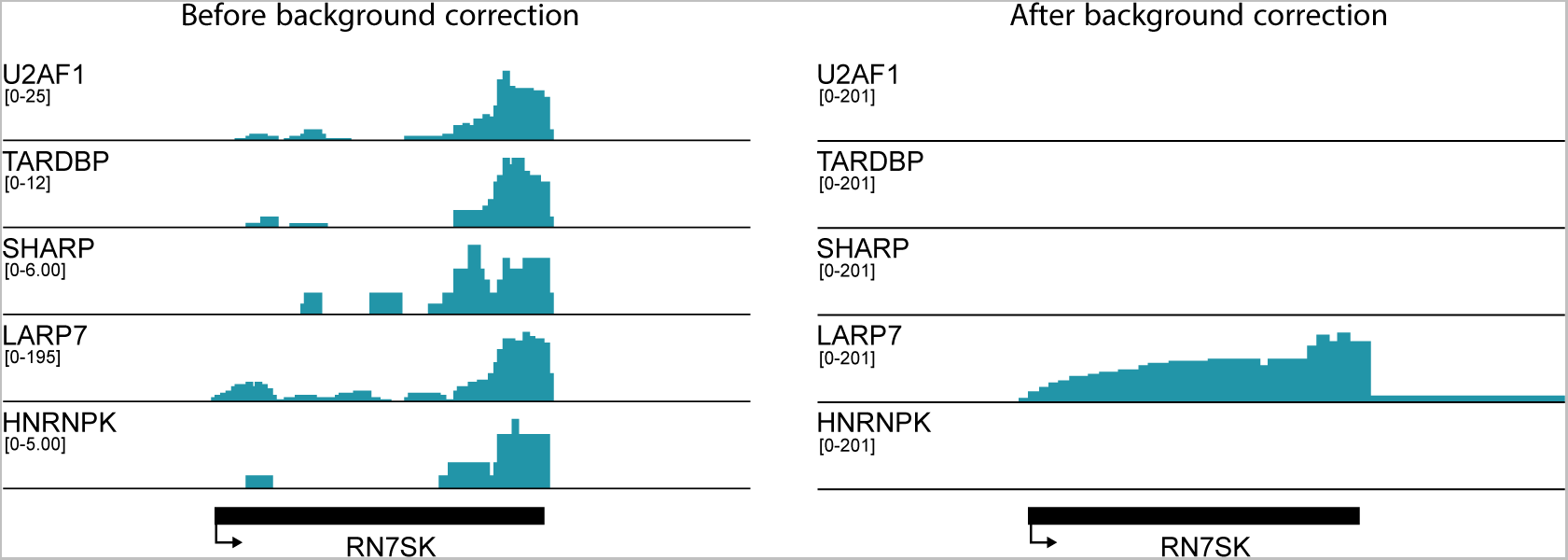
Background correction. An example of our background correction method that utilizes the total read coverage across all proteins to normalize each individual protein. Shown are example tracks on RN7SK before and after background correction. Left: Raw alignment data for the entire pooled dataset (top track) and for representative antibodies against U2AF1, TARDBP, SHARP, LARP7 and HNRNPK on *RN7SK*. Right: Background corrected data for the same set of antibodies. Signal that was not antibody-specific has been normalized out. The reads in the right are binned in 5 nucleotide windows. *RN7SK* is known to be bound by LARP7Ref.51.

**Supplemental Figure 6:**
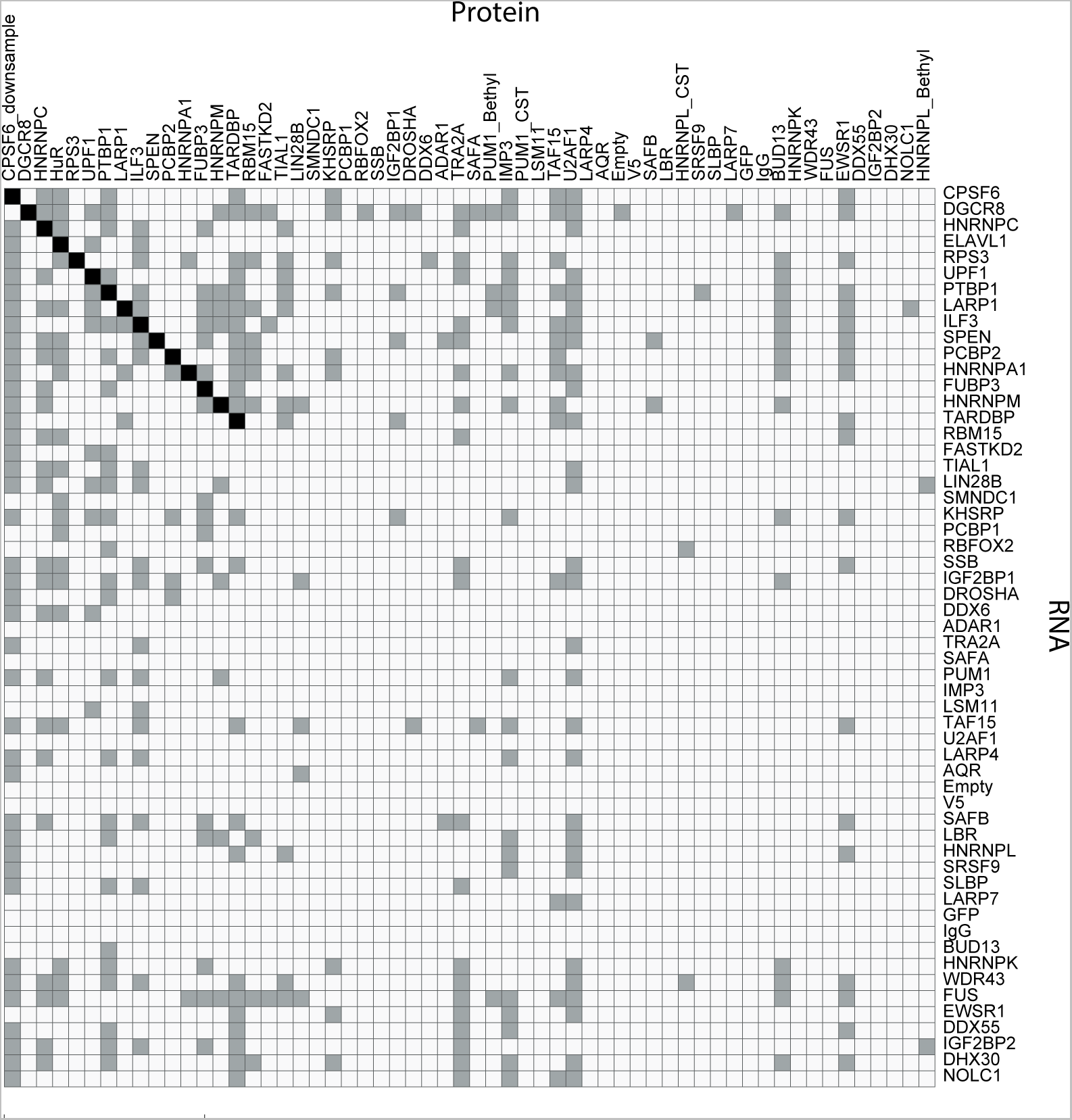
Autoregulatory binding between RBPs targeted by SPIDR and their RNAs. Auto-regulatory binding matrix with protein (x-axis) binding to each mRNA (y-axis) shown. Each target protein included in SPIDR performed in K562 cells marked by whether it has significantly enriched binding within its own RNA, or in any of the other SPIDR target RNAs. Proteins that bind their own RNA are marked in black, instances of binding to genes of other SPIDR targets are marked in gray.

**Supplemental Figure 7:**
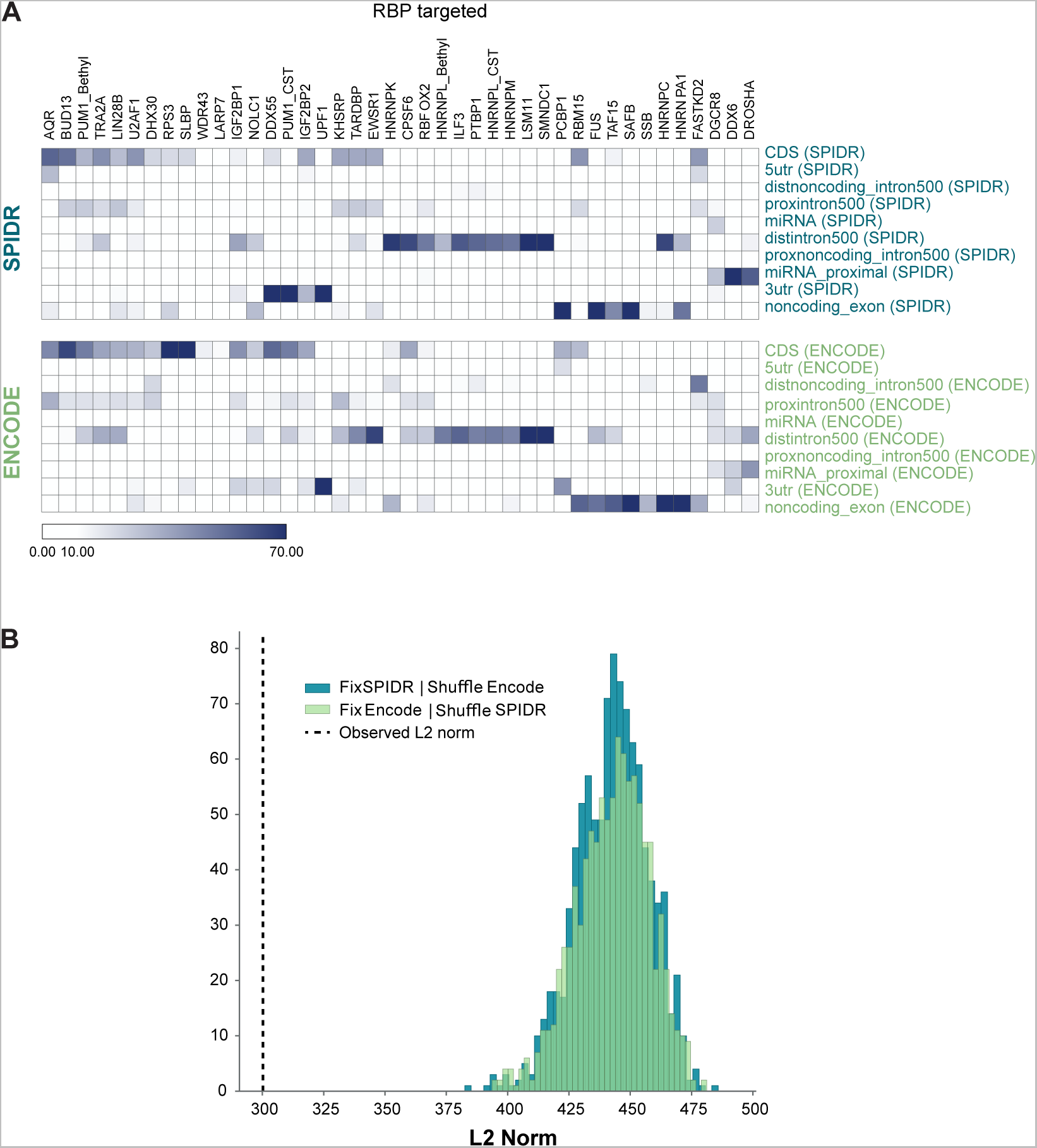
Global comparison of annotations (intron, exon, etc) of binding sites per RBP as called by ENCODE versus SPIDR. **(A)** Heatmaps showing the percentage of significant binding sites in each of the annotation categories for SPIDR performed in K562 cells and ENCODE (see **Methods** for details). **(B**) Quantitative assessment of the similarity of heatmaps between SPIDR and ENCODE. The Euclidean distance (L2 norm) between the ENCODE and SPIDR percentage tables/heatmaps was calculated. The calculated distance is indicated by the dashed line. The statistical significance was calculated by randomly shuffling the columns of either the SPIDR percentage table and keeping the original ENCODE table or vice versa, meaning shuffling the columns of the ENCODE table and keeping the original SPIDR table. This was done 1000 times in each direction and every time the Euclidean distance was calculated. The values are represented by the two histograms. The Euclidean distance of all of the randomly shuffled 2000 comparison was always larger than of the true pair, which shows that the two original annotation tables from SPIDR and ENCODE are highly significantly similar (p-value < 0.0005).

**Supplemental Table 1:** Overview of the SPIDR experiment in K562 cells. The protein targets are listed, as well as the vendors and product numbers of the corresponding antibodies.

| Target RBP | Antibody Vendor | Antibody Catalog Number |
| --- | --- | --- |
| HNRNPM | Santa Cruz | sc-20001 |
| HNRNPC | Abcam | ab10294 |
| HNRNPL Bethyl | Bethyl | A303-896A |
| RBM15 | Bethyl | A300-821A |
| DHX30 | Bethyl | A302-218A |
| SAFB | Invitrogen | MA1-91526 |
| PCBP1 | Abcam | 168378 |
| V5 | Invitrogen | MA5-15253 |
| SRSF9 | Abcam | ab74782 |
| ADAR1 | CST | D7E2M |
| HuR | CST | D9W7E |
| AQR | Bethyl | A302-547A |
| FUBP3 | Bethyl | A303-012A |
| ILF3 | Bethyl | A303-651A |
| LARP4 | Bethyl | A303-723A |
| BUD13 | Bethyl | A303-320A |
| DDX52 | Bethyl | A303-054A |
| DDX55 | Bethyl | A303-027A |
| DDX6 | Bethyl | A300-460A |
| DGCR8 | Bethyl | A302-468A |
| DROSHA | Bethyl | A301-886A |
| EWSR1 | Bethyl | A300-417A |
| FASTKD2 | Bethyl | A303-788A |
| LARP7 | Bethyl | A303-723A |
| TRA2A | Bethyl | A303-779A |
| U2AF1 | Bethyl | A302-079A |
| UPF1 | Bethyl | A301-902A |
| LIN28B | Bethyl | A303-588A |
| NOLC1 | Bethyl | A302-184A |
| PUM1 Bethyl | Bethyl | A302-576A |
| RBFOX2 | Bethyl | A300-864A |
| RPS3 | Bethyl | A303-840A |
| SHARP | Bethyl | A301-119A |
| TAF15 | Bethyl | A300-308A |
| TARDBP | Bethyl | A303-223A |
| CPSF6 | CST | 92879S |
| IGF2BP1 (IMP1) | MBL International | RN007P |
| KHSRP | Abcam | ab150393 |
| LARP1 | Bethyl | A303-900A |
| FUS | CST | 67840S |
| HNRNPL CST | CST | 65043S |
| IMP3 | CST | 57145S |
| PTBP1 | CST | 57246S |
| PUM1 CST | CST | 12322S |
| TIAL1 | CST | 8509S |
| HNRNPA1 | Abcam | ab4791 |
| HNRNPK | MBL International | RNP019 |
| PCBP2 | MBL International | RN025P |
| WDR43 | Bethyl | A302-478A |
| LSM11 | Bethyl | A303-709A |
| Empty | n/a | n/a |
| SSB | MBL International | RN074PW |
| SLBP | MBL International | RN045P |
| SMNDC1 | MBL International | RN078PW |
| GFP | Living Colors | JL-8 |
| IgG | Abcam | ab172730 |
| IGF2BP1 (IMP2) | MBL International | RN008P |
| LBR | Abcam | ab122919 |
| HNRNPU (SAF-A) | Abcam | ab20666 |

**Supplemental Table 2:** Overview of the SPIDR experiment HEK293T cells treated with Torin or Control (solvent only). The protein targets are listed, as well as the vendors and product numbers of the corresponding antibodies.

| Target RBP | Antibody Vendor | Antibody Catalog Number |
| --- | --- | --- |
| AQR | Bethyl | A302-547A |
| SAF-A | Abcam | ab20666 |
| HNRNPM3/4 | Santa Cruz | sc-20001 |
| BUD13 | Bethyl | A303-320A |
| DDX52 | Bethyl | A303-054A |
| DDX55 | Bethyl | A303-027A |
| DDX6 | Bethyl | A300-460A |
| DGCR8 | Bethyl | A302-468A |
| DROSHA | Bethyl | A301-886A |
| EWSR1 | Bethyl | A300-417A |
| V5 | Invitrogen | MA5-15253 |
| FASTKD2 | Bethyl | A303-788A |
| SHARP | Bethyl | A301-119A |
| TAF15 | Bethyl | A300-308A |
| FUBP3 | Bethyl | A303-900A |
| ILF3 | Bethyl | A303-651A |
| LARP4 | Bethyl | A303-900A |
| LARP7 | Bethyl | A303-723A |
| LIN28B | Bethyl | A303-588A |
| NOLC1 | Bethyl | A302-184A |
| RBFOX2 | Bethyl | A300-864A |
| RPS3 | Bethyl | A303-840A |
| TRA2A | Bethyl | A303-779A |
| HNRNPL | Bethyl | A303-896A |
| IMP3 | CST | 57145S |
| PTBP1 | CST | 57246S |
| U2AF1 | Bethyl | A302-079A |
| UPF1 | Bethyl | A301-902A |
| EIF4G1 | CST | 2858S |
| RPS2 | Bethyl | A303-794A |
| RBM15 | Bethyl | A300-821A |
| CPSF6 | CST | 92879S |
| HNRNPC | Abcam | ab10294 |
| PUM1 | Bethyl | A302-576A |
| SLBP | MBL International | RN045P |
| SMNDC1 | MBL International | RN078PW |
| GFP | Living Colors | JL-8 |
| TIAL-1 | CST | 8509S |
| HNRNPK | MBL International | RNP019 |
| IMP1 | MBL International | RN007P |
| KHSRP | Abcam | ab150393 |
| LARP1 | Bethyl | A303-900A |
| SSB | MBL International | RN074PW |
| IgG | Abcam | ab172730 |
| XRN1 | CST | 70205S |
| RPS6 | CST | 2217S |
| Empty | n/a | n/a |
| IMP2 | MBL International | RN008P |
| SAFB | Invitrogen | MA1-91526 |
| LBR | Abcam | ab122919 |
| PCBP1 | Abcam | 168378 |
| 4EBP1 | CST | 9644S |
| EIF4E | CST | 2067S |
| EIF4A | CST | 2013S |

## MATERIALS AND METHODS

### Experimental conditions

#### Cell culture

K562 cells (ATCC, CCL-243) and HEK293T cells (ATCC, CRL-3216) were purchased from ATCC and cultured under standard conditions. K562 cells were cultured in K562 media consisting of 1X DMEM (Gibco), 1 mM Sodium Pyruvate (Gibco), 2 mM L-Glutamine (Gibco), 1X FBS (Seradigm), 100 U/mL Penicillin-Streptomycin (Life Technologies). HEK293T cells were cultured in HEK293T media consisting of 1X DMEM media (Gibco), 1 mM MEM non-essential amino acids (Gibco), 1 mM Sodium Pyruvate (Gibco), 2 mM L-Glutamine (Gibco), 1X FBS (Seradigm).

#### UV-crosslinking

Crosslinking was performed as previously described^23^. Briefly, K562 cells were washed once with 1X PBS and diluted to a density of ~10 million cells/mL in 1X PBS for plating onto culture dishes. HEK293T cells were washed once with 1X PBS and crosslinked directly on culture dishes. RNA-protein interactions were crosslinked on ice using 0.25 J cm^−2^ (UV 2.5k) of UV at 254 nm in a Spectrolinker UV Crosslinker. Cells were then scraped from culture dishes, washed once with 1X PBS, pelleted by centrifugation at 330 x g for 3 minutes, and flash-frozen in liquid nitrogen for storage at −80°C.

#### Torin-1 treatment

HEK293T cells were treated at a final concentration of 250 nM Torin-1 (Cell Signaling Technology, #14379) in standard HEK293T media for 18 hours prior to UV-crosslinking and harvesting.

#### Bead biotinylation

The bead labeling strategy was adapted from ChIP DIP, a Guttman lab protocol used for multiplexed mapping of hundreds of proteins the DNA (https://guttmanlab.caltech.edu/technologies/). Specifically, 1 mL of Protein G Dynabeads (ThermoFisher, #10003D) were washed once with 1X PBST (1X PBS + 0.1% Tween-20) and resuspended in 1mL PBST. Beads were then incubated with 20 μL of 5 mM EZ-Link Sulfo-NHS-Biotin (Thermo, #21217) on a HulaMixer for 30 minutes at room temperature. Following NHS reaction, beads were placed on a magnet and 500 μL of buffer was removed and replaced with 500 μL of 1M Tris pH 7.4 to quench the reaction for an additional 30 minutes at room temperature. Beads were then washed twice with 1 mL PBST and resuspended in their original storage buffer until use.

#### Labeling biotinylated beads with oligonucleotide tags

Unique biotinylated oligonucleotides were first coupled to streptavidin (BioLegend, #280302) in a 96-well PCR plate. In each well, 20 μL of 10 μM oligo was added to 75 μL 1X PBS and 5 μL 1 mg/mL streptavidin. The 96-well plate was then incubated with shaking at 1600 rpm on a ThermoMixer for 30 minutes at room temperature. Each well was then diluted 1:4 in 1X PBS for a final concentration of 227 nM.

For each experiment, the appropriate amount of biotinylated Protein G beads (10 μL beads per capture antibody) was washed once in 1X PBST. Beads were then resuspended in oligo binding buffer (0.5X PBST, 5 mM Tris pH 8.0, 0.5 mM EDTA, 1M NaCl). 200 μL of the bead suspension was aliquoted into individual wells of a 96-well plate, followed by addition of 4 μL of 227nM streptavidin-coupled oligo to each well. The 96-well plate was then incubated with shaking at 1200 rpm on a ThermoMixer for 30 minutes at room temperature. Beads were then washed twice with M2 buffer (20 mM Tris 7.5, 50 mM NaCl, 0.2% Triton X-100, 0.2% Na-Deoxycholate, 0.2% NP-40), twice with 1X PBST, and resuspended in 200 μL of 1X PBST.

#### Binding antibody to labeled Protein G beads

2.5 μg of each capture antibody was added to each well of the 96-well plate containing labeled beads in 1X PBST. The plate was incubated with shaking at 1200 rpm on a ThermoMixer for 30 minutes at room temperature. After incubation, beads were washed twice with 1X PBST + 2 mM biotin (Sigma, #B4639-5G), resuspended in 200 μL of 1x PBST + 2mM biotin, and left shaking at 1200 rpm for 10 minutes at room temperature. All wells containing beads were then pooled together and washed twice with 1 mL 1X PBST + 2 mM biotin. At this stage, each bead in the bead pool contains a single type of capture antibody with a corresponding unique oligonucleotide tag.

#### Pooled immunoprecipitation

For each experiment, 10 million cells were lysed in 1 mL RIPA buffer (50mM HEPES pH 7.4, 100mM NaCl, 1% NP-40, 0.5% Na-Deoxycholate, 0.1% SDS) supplemented with 20 μL Protease Inhibitor Cocktail (Sigma, #P8340-5mL), 10 μL of Turbo DNase (Invitrogen, #AM2238), 1X Manganese/Calcium mix (2.5 mM MnCl_2_, 0.5 mM CaCl_2_), and 5 μL of RiboLock RNase Inhibitor (Thermo Fisher, #EO0382)). Samples were incubated on ice for 10 minutes to allow lysis to proceed. After lysis, cells were sonicated at 3-4 W of power for 3 minutes (pulses 0.7 s on, 3.3 s off) using the Branson sonicator and then incubated at 37°C for 10 minutes to allow for DNase digestion. DNase reaction was quenched with addition of 0.25 M EDTA/EGTA mix for a final concentration of 10 mM EDTA/ EGTA. RNase If (NEB, #M0243L) was then added at a 1:500 dilution and samples were incubated at 37°C for 10 minutes to allow partial fragmentation of RNA to obtain RNAs of approximately ~300-400 bp in length. RNase reaction was quenched with addition of 500 μL ice cold RIPA buffer supplemented with 20 μL Protease Inhibitor Cocktail and 5 μL of RiboLock RNase Inhibitor, followed by incubation on ice for 3 minutes. Lysates were then cleared by centrifugation at 15000 x g at 4°C for 2 minutes. The supernatant was transferred to new tubes and diluted in additional RIPA buffer such that the final volume corresponded to 1 mL lysate for every 100 μL of Protein G beads used. Lysate was then combined with the labeled antibody-bead pool and 1 M biotin was added to a final concentration of 10 mM as to quench any disassociated streptavidin-coupled oligos. Beads were left rotating overnight at 4°C on a HulaMixer. Following immunoprecipitation, beads were washed twice with RIPA buffer, twice with high salt wash buffer (50 mM HEPES pH 7.4, 1 M NaCl, 1% NP-40, 0.5% Na-Deoxycholate, 0.1% SDS), and twice with Tween buffer (50 mM HEPES pH 7.4, 0.1% Tween-20).

#### Ligation of the RNA Phosphate Modified (“RPM”) tag

After immunoprecipitation, 3’ ends of RNA were modified to have 3’ OH groups compatible for ligation using T4 Polynucleotide Kinase (NEB, #M0201L). Beads were incubated at 37°C for 10 minutes with shaking at 1200 rpm on a ThermoMixer. Following end repair, beads were buffer exchanged by washing twice with high salt wash buffer and twice with Tween buffer. RNA is subsequently ligated with an “RNA Phosphate Modified” (RPM) adaptor (Quinodoz et al 2021) using High ConcentrationT4 RNA Ligase I (NEB, M0437M). Beads were incubated at 24°C for 1 hour 15 minutes with shaking at 1400 rpm, followed by three washes in Tween buffer. After RPM ligation, RNA was converted to cDNA using SuperScript III (Invitrogen, #18080093) at 42°C for 20 minutes using the “RPM Bottom” RT primer to facilitate on-bead library construction and a 5’ sticky end to ligate tags during split-and-pool barcoding. Excess primer is digested with Exonuclease I (NEB, #M0293L) at 37°C for 15 minutes.

#### Split-and-pool barcoding to identify RNA-protein interactions

Split-and-pool barcoding was performed as previously described^31^ with minor modifications. Specifically, beads were split-and-pool ligated over ≥ 6 rounds with a set of “Odd,” “Even,” and “Terminal” tags. The number of barcoding rounds performed for each SPIDR experiment was determined based on the complexity of the given bead pool. All split-and-pool ligation steps were performed for 5 minutes at room temperature and supplemented with 2 mM biotin and 1:40 RiboLock RNase Inhibitor to prevent RNA degradation. We ensured that virtually all barcode clusters (>95%) represented molecules belonging to unique, individual beads.

Compared to previously published approaches, we reduced the number of barcodes per round, but increased the rounds of split and pool barcoding as we optimized the ligation step. Therefore, the barcoding procedure was significantly simplified in contrast to previous versions. For example, for the K562 cells pooled experiment, 6 rounds of 24 barcodes were used for combinatorial barcoding (with a scheme of Odd, Even, Odd, Even, Odd, Terminal tag). For the HEK293T cells mTOR inhibition experiment, 6 rounds of 36 barcodes were used for combinatorial barcoding to achieve sufficient barcode complexity. Of the 36 barcodes used in round one of the ligations, 18 were used to label the control condition and the remaining 18 were used to label the torin treated condition. The samples were then pooled together for the remaining 5 rounds of ligation.

#### Library preparation

After split-and-pool barcoding, beads were aliquoted into 5% aliquots for library preparation and sequencing. RNA in each aliquot was degraded by incubating with RNase H (NEB, #M0297L) and RNase cocktail (Invitrogen, #AM2286) at 37°C for 20 minutes. 3’ ends of the resulting cDNA were ligated to attach dsDNA oligos containing library amplification sequences using a “splint” ligation as previously described (Quinodoz et al 2021)^31^. The “splint” ligation reaction was performed with 1X Instant Sticky End Master Mix (NEB #M0370) at 24°C for 1 hour with shaking at 1400 rpm on a ThermoMixer. Barcoded cDNA and biotinylated oligo tags were then eluted from beads by boiling in NLS elution buffer (20 mM Tris-HCl pH 7.5, 10 mM EDTA, 2% N-lauroylsarcosine, 2.5 mM TCEP) for 6 minutes at 91°C, with shaking at 1350 rpm.

Biotinylated oligo tags were first captured by diluting the eluant in 1X oligo binding buffer (0.5X PBST, 5 mM Tris pH 8.0, 0.5 mM EDTA, 1M NaCl) and subsequently binding to MyOne Streptavidin C1 Dynabeads (Invitrogen, #65001) at room temperature for 30 minutes. Beads were placed on a magnet and the supernatant, containing cDNA, was moved to a separate tube. Biotinylated oligo tags were amplified on-bead using 2X Q5 Hot-Start Mastermix (NEB #M0494) with primers that add the indexed full Illumina adaptor sequences.

To isolate barcoded cDNA, the supernatant was first incubated with a biotinylated antisense ssDNA (“anti-RPM”) probe that hybridizes to the junction between the reverse transcription primer and splint sequences to reduce empty insertion products. This mixture was then bound to MyOne Streptavidin C1 Dynabeads at room temperature for 30 minutes. Beads were placed on a magnet and the supernatant, containing the remaining cDNA products, was cleaned up on Silane beads (Invitrogen, #37002D) as previously described^83^. Finally, cDNA was amplified using 2X Q5 Hot-Start Mastermix (NEB #M0494) with primers that add the indexed full Illumina adaptor sequences.

After amplification, libraries were cleaned up using 1X SPRI (AMPure XP), size-selected on a 2% agarose gel, and cut at either ~300 nt (barcoded oligo tag) or between 300-1000 nt (barcoded cDNA). Libraries were subsequently purified with Zymoclean Gel DNA Recovery Kit (Zymo Research, #4007).

#### Sequencing

Paired-end sequencing was performed on either an Illumina NovaSeq 6000 (S4 flowcell), NextSeq 550, or NextSeq 2000 with read lengths ≥ 100 x 200 nucleotides. For the K562 data, 37 SPIDR aliquots were generated and sequenced from two technical replicate experiments. The two experiments were generated using the same batch of UV-crosslinked lysate processed on the same day. For the HEK293T data, 9 SPIDR aliquots were generated from a single technical replicate. Each SPIDR library corresponds to a distinct aliquot that was separately amplified with different indexed primers, providing an additional round of barcoding as previously described^31^. Minimum required sequencing depth for each experiment was determined by the estimated number of beads and unique molecules in each aliquot. For oligo tag libraries, each library was sequenced to a depth of observing ~5 unique oligo tags per bead on average. For cDNA libraries, each library was sequenced with at least 2x coverage of the total estimated library complexity.

### Analysis and processing pipeline

#### Read processing and alignment

Paired-end RNA sequencing reads were trimmed to remove adaptor sequences using Trim Galore! v0.6.2 and assessed with FastQC v0.11.8. Subsequently, the RPM (ATCAGCACTTA) sequence was trimmed using Cutadapt v3.4 from both 5’ and 3’ read ends. The barcodes of trimmed reads were identified with Barcode ID v1.2.0 (<u>https://</u> <u>github.com/GuttmanLab/sprite2.0-pipeline</u>) and the ligation efficiency was assessed. Reads with or without an RPM sequence were split into two separate files to process RNA and oligo tag reads individually downstream, respectively.

RNA read pairs were then aligned to a combined genome reference containing the sequences of repetitive and structural RNAs (ribosomal RNAs, snRNAs, snoRNAs, 45S pre-rRNAs, tRNAs) using Bowtie2. The remaining reads were then aligned to the human (hg38) genome using STAR aligner. Only reads that mapped uniquely to the genome were kept for further analysis.

#### Barcode matching and filtering

Mapped RNA and oligo tag reads were merged, and a cluster file was generated for all downstream analysis as previously described. MultiQC v1.6 was used to aggregate all reports. To unambiguously exclude ligation events that could not have occurred sequentially, we utilized unique sets of barcodes for each round of split-and-pool. All clusters containing barcode strings that were out-of-order or contained identical repeats of barcodes were filtered from the merged cluster file. To determine the amount of unique oligo tags present in each cluster, sequences sharing the same Unique Molecular Identifier (UMI) were removed and the remaining occurrences were counted. To remove PCR duplication events within the RNA library, sequences sharing identical start and stop genomic positions were removed.

#### Splitting alignment files by protein identity

Barcode strings from filtered cluster files were then used to assign protein identities to the alignment file containing all mapped RNA reads. Because each cluster represents an individual bead, the frequency of oligo tags (each representing unique protein type) was used to determine protein assignments. Specifically, for each cluster we required ≥ 3 observed oligo tags and that the most common protein type represented ≥ 80% of all observed tags. RNA reads were then split into separate alignment files by barcode strings corresponding to protein type.

#### Background correction and peak calling

In order to determine what portion of the observed signal is specific to a particular capture antibody, rather than common pileups regardless of the protein captured, we normalized coverage for each protein relative to the coverage detected for all other proteins. Specifically, for a protein of interest, we computed the number of reads that were mapped to that protein. We then randomly downsampled all reads not assigned to that protein such that it had a comparable number of reads as the protein of interest. To measure the expected variance in the control sample, we repeated this downsampling procedure at least 100 independent times. We then computed read counts per window across the transcriptome (either 10nts or 100nts) for the protein of interest and each of the randomized control samples. We computed a normalized enrichment as the number of observed reads within the window (observed) divided by the average of the read counts overt that window across the >=100 permutations (expected). To assess the significance of this enrichment score, we measured how often the observed score was seen in the >=100 permutations. A p-value was assigned as the number of random scores greater than or equal to the observed scores divided by the number of random permutations used (we included the actual observed score in the numerator and denominator). All windows that had at least 10 observed reads and a p-value less than 0.05 were considered significantly enriched.

#### Peak annotation

Enriched windows were first filtered to only include regions resulting from reads that could be uniquely mapped in the second STAR alignment, and then poor alignments to rRNA regions (chr21: 88206400-8449330) were removed. These filtered peaks were then annotated based on overlap with GENCODE v41 transcripts. In the case of overlapping annotations, the final assigned annotation was chosen based on the following priority list: miRNA, CDS, 5’UTR, 3’UTR, proximal intron (within 500 nt of the splice site region), distal intron (further than 500 nt of the splice site region), non-coding exon, and finally non-coding intron. Windows for which the primary gene annotation was a miRNA host gene were marked as miRNA proximal.

### SPIDR comparison to ENCODE

#### ENCODE datasets

43 of the proteins included in SPIDR also had a matched K562 ENCODE eCLIP experiment with paired-end sequencing data. The raw FASTQ files for these datasets were downloaded from the ENCODE website (https://www.encodeproject.org/) and aligned to the genome using the same parameters as in the SPIDR dataset.

For comparison of matched SPIDR and ENCODE datasets, the larger of the pair of alignment files was downsampled to the depth of the smaller alignment file. Windows of enrichment in ENCODE datasets were then determined using the same background correction strategy and thresholding as in SPIDR (minimum read count of 10, p-value < 0.05). As was done in the SPIDR data, all ENCODE datasets were used as negative controls for one for one another when determining background correction factors and calling windows of enrichment.

#### Motif enrichment analysis

Filtered SPIDR peaks were used to subset the corresponding SPIDR alignment files, such that only reads that fell within enriched windows were kept. These reads were then used as input for *de novo* motif analysis by HOMER (http://homer.ucsd.edu/homer/). Motifs with a reported p-value < 10^−40^ were considered significant.

#### Comparison of bound RNA features

Enriched windows for both SPIDR and ENCODE, as determined using the SPIDR workflow of background correction and thresholding, were annotated based on overlap with GENCODE v41 transcripts. Peaks annotated as intergenic were removed, and then both the SPIDR and ENCODE datasets were filtered to include only proteins that had greater than 100 peaks.

The likelihood of seeing a similarity between SPIDR and ENCODE in the region annotations is visualized by comparing the observed values to randomly shuffled values. The inputs for this method are two matrices, one for SPIDR and one for ENCODE, with the percentage of annotations observed for a given region type for a given RBP. Shuffling is performed by randomly switching percentages across RBPs, keeping the relative values between regions constant. This can be thought of as randomly shuffling the columns of one of the input matrices. A distance is calculated by flattening the two input matrices into vectors, taking the difference between the two vectors, and calculating an L2-norm on that difference. In Supplemental Figure 7 the histogram of L2-norms shows the distribution we would expect if RBPs had no effect on the L2-norm between SPIDR and ENCODE. The dashed vertical line represents the L2-norm when the input matrices were flattened but not shuffled.

The basic algorithm is as follows:

Calculate the true L2-norm between SPIDR and ENCODE

Keeping SPIDR constant, randomly switch probabilities between RBPs while keeping the percentages within an RBP the same for ENCODE

Repeat step 2, shuffling SPIDR and keeping ENCODE constant

Repeat steps 2 and 3 for 1000 samples

#### Single nucleotide resolution analysis

We computed the frequency of reads ending at the 3’ end of the cDNA. We computed enrichment for each of these counts by randomly downsampling all reads not assigned to the specific protein and computing the same 3’ end coverage. Enrichments and p-values were computed as described above and as previously reported in (Banerjee et al. 2020)^84^.

#### mTOR analysis

Background corrected bedgraphs were generated from control and +Torin conditions for each RBP in each condition. These bedgraph values were then mapped on to Refseq genes using the bedtools map command (arguments: -c 4 –o absmax). Where multiple isoforms were present for the same gene, the isoform with the highest map count was used. To normalize for possible detection bias due to fewer antibody beads in one condition versus the other we adjusted the map value by the ratio of antibody beads as determined by number of bead clusters corresponding to each antibody in each respective condition. Number of antibody (bead clusters) were defined and calculated using the same values used to generate the split bam files for each protein (options: minimum number of oligos=3,fraction unique=0.8, max number of RNAs in clusters=100). The ratio of cluster-corrected values for each gene across the two conditions was then compared per gene and separated based on TOP score. Published TOP scores^60^ were used to generate categories for violin plots.

For the protein changes CDF plots, we first selected for the 2000 highest expressed genes based on previous RNA-seq data^84^. Input TPM values for HEK293 cells were taken from input CLAP (sub_input.merged.bam) data from HEK293T cell in (Banerjee et al. 2020)^84^.The input samples were downsampled to 20M reads prior to TPM calculation. Featurecounts was used to calculate read overlaps with hg38 protein coding refseq genes and further converted to TPM values. The top 2000 expressed genes (based on HEK293 input TPM) were used to plot the average protein log2 fold changes (Torin versus control) vs TOP score. Published TOP scores^60^ were used to plot CDF values.

### Mass spectrometry

#### Multiplexed Immunopurification (IP) for mass spectrometry

10 million K562 cells were lysed in 4mL of RIPA on ice for 10 minutes. The lysate was clarified by centrifugation at 15000g for 2 minutes, and then split in half for either the pooled IP with 39 antibodies or the negative control IP with an anti-V5 antibody. Each half of the lysate was combined with 10ug total antibody (0.25ug per each antibody for the pooled IP) and 100uL of Protein G beads and left rotating at 4C overnight. The beads were then washed twice with RIPA, twice with High Salt Wash Buffer, twice with Clap-Tween, and finally three times with Mass Spec IP Wash Buffer (150mM NaCl, 50mM Tris-HCl pH 7.5, 5% Glycerol). Each sample was then reduced, alkylated, Trypsin digested, and desalted as described in (Parnas et al, 2015)^85^. Peptides were reconstituted in 12uL 3% acetonitrile/0.1% formic acid.

#### mTOR proteomics

5 million cells each of control and 250nM Torin-1 treated HEK cells were lysed in 250uL Mass Spec Lysis Buffer (8M urea, 75mM NaCL, 50mM Tris pH 8.0, 1mM EDTA) for 30min at room temperature. Samples were then clarified by centrifugation at 23000g for 5 minutes, and the protein content in the supernatant was measured by BCA assay (ThermoFisher, #PI23227). 40ug of protein for each sample was reduced with 5mM final dithriothreitol (DTT) for 45 minutes at room temperature and subsequently alkylated with 10mM final iodoacetamide (IAA) for 45 minutes in the dark at room temperature. 50mM Tris (pH 8.0) was then added to each sample such that the final concentration of urea was less than 2M. Samples were digested overnight with 0.4ug Trypsin (Promega, #V5113) for a 1:100 enzyme to protein ratio. Peptides were desalted on C18 StageTips according to (Rappsilber et al., 2007)^86^.

#### LC-MS/MS

LC-MS/MS analysis was performed on a Q-Exactive HF. 5uL of total peptides were analyzed on a Waters M-Class UPLC using a C18 25cm Thermo EASY-Spray column (2um, 100A, 75um x 25cm) or IonOpticks Aurora ultimate column (1.7um, 75um x 25cm) coupled to a benchtop ThermoFisher Scientific Orbitrap Q Exactive HF mass spectrometer. Peptides were separated at a flow rate of 400 nL/min with a linear 95 min gradient from 5% to 22% solvent B (100% acetonitrile, 0.1% formic acid), followed by a linear 30 min gradient from 22 to 90% solvent B. Each sample was run for 160 min, including sample loading and column equilibration times. Data was acquired using Xcalibur 4.1 software.

The IP samples were measured in a Data Dependent Acquisition (DDA) mode. MS1 Spectra were measured with a resolution of 120,000, an AGC target of 3e6 and a mass range from 300 to 1800 m/z. Up to 12 MS2 spectra per duty cycle were triggered at a resolution of 15,000, an AGC target of 1e5, an isolation window of 1.6 m/z and a normalized collision energy of 28.

The Torin treated and control total lysate samples were measured in a Data Independent Acquisition (DIA) mode. MS1 Spectra were measured with a resolution of 120,000, an AGC target of 5e6 and a mass range from 350 to 1650 m/z. 47 isolation windows of 28 m/z were measured at a resolution of 30,000, an AGC target of 3e6, normalized collision energies of 22.5, 25, 27.5, and a fixed first mass of 200 m/z.

#### Database searching of the proteomics raw files

Proteomics raw files were analyzed using the directDIA method on SpectroNaut v16.0 for DIA runs or SpectroMine (3.2.220222.52329) for DDA runs (Biognosys) using a human UniProt database (Homo sapiens, UP000005640), under BSG factory settings, with automatic cross-run median normalization and imputation. Protein group data were exported for subsequent analysis.

